# Targeting of the barley cell-surface receptor SRF3 by the *Blumeria hordei* effector AVRA13 overlaps with AVRA13 recognition by MLA and the induction of NLR-mediated cell death

**DOI:** 10.64898/2026.01.08.698370

**Authors:** Wei Shi, Merle Bilstein-Schloemer, Sara C. Stolze, Matthieu P. Platre, Pichaporn Chuenban, Sophie C. Sent, Ishani S. Das, Maike Hofmann, Götz Hensel, Gunther Doehlemann, Hirofumi Nakagami, Isabel M.L. Saur

## Abstract

Pathogens secrete effector proteins to promote virulence. Despite their recognition by barley *Mla* resistance genes, the structurally-related *Blumeria hordei* (*Bh*) *AVR*_a_ effectors are maintained in the *Bh* genome, suggesting virulence functions critical for fungal pathogenicity. Using proximity-dependent protein labelling in transgenic barley, we detected distinct host protein interactomes for five AVR_A_s despite their structural homology and convergence on MLAs. We report the specific interaction of the highly conserved AVR_A13_ effector with the barley cell-surface receptor SRF3. AVR_A13_ disrupts HvSRF3-HvBAK1 interaction and alters HvSRF3 plasma membrane levels. AVR_A13_ can also bind *Arabidopsis thaliana* SRF3 and *A. thaliana AVR_a13_*-lines mimic *srf3* mutants in iron signaling. In barley, *srf3* mutants and *AVR_a13_* expression desensitize iron-induced restriction of *Bh* growth, suggesting that AVR_A13_ facilitates fungal proliferation by manipulating SRF3-mediated changes to iron homeostasis. Together, these findings identify SRF3 as molecular link between pathogen virulence and iron-related processes. In addition, our data demonstrates that MLA specifically detects the residues that underlie *Bh* effector neo-functionalization and intrinsic AVR_A_ virulence functions.

## Introduction

### Plant immune receptor-mediated non-self-recognition and pathogen virulence effectors

To defend against pathogenic microbes, plants trigger immune responses upon detecting conserved microbial patterns, such as fungal chitin or bacterial flagellin, *via* cell-surface receptor-like kinases (RLKs), many of which use the co-receptor BAK1 (BRASSINOSTEROID INSENSITIVE 1-ASSOCIATED KINASE 1). This response is known as pattern-triggered immunity (PTI)^1,2^. Plant-adapted microbial invaders secrete effectors to suppress host immunity and acquire nutrients, which is the intrinsic effector activity^3,4^. Host resistance to adapted pathogens is mediated by plant *Resistance* (*R*) genes, that frequently encode intracellular nucleotide-binding oligomerization domain-like receptors (NLRs), which detect pathogen effectors^5^. Effector-mediated activation of NLRs is commonly associated with a localized cell death response^6^ that eliminates the host niche of biotrophic pathogens as they feed of living host cells^6^. Pathogen effectors recognized by R proteins are historically known as ‘avirulence effectors’ (AVRs) and this activity can be considered the effector’s avirulence function^2^. Recognition of AVRs by direct interaction with an immune receptor is typically associated with rapid evasion of NLR recognition by accumulation of effector gene polymorphisms in pathogen populations^6^.

The obligate biotrophic powdery mildew fungi infect a wide range of plants including economically relevant crops^7^. The barley powdery mildew fungus *Blumeria hordei* (*Bh*) harbors hundreds of candidate secreted effector protein (CSEP)-coding genes^8,9^. The large number of CSEPs paired with the difficulties in genetically modifying obligate biotrophs makes it highly challenging to experimentally identify the effectors’ virulence function crucial for barley host infection. This constitutes a major knowledge gap in the molecular understanding of the establishment of an obligate biotrophic lifestyl^10^. The diversified *Mildew locus a* encodes sequence-related NLRs (MLAs) with recognition specificities towards effectors derived from multiple unrelated fungal phytopathogens, among them the *Bh* AVR_A_ effectors^11–13^. To date, seven *AVR_a_* effectors have been isolated^14–16^. All belong to different subclasses of the expanded *Bh* RALPH (RNase-Like Proteins expressed in Haustoria) effector family^14,16–18^. The recognition of AVR_A_ effectors by intracellular MLA NLRs demonstrates AVR_A_ translocation into the host cell and their predominant expression upon haustorium formation underscores the secretion *via* the haustorium, a specialized fungal infection structure which is the primary site for the exchange of molecules between host and pathogen^10,17,19–21^. Like most RALPH members, AVR_A_ effectors are pseudo-RNases lacking catalytic residues, which suggests that they evolved from an ancient ribonuclease with neo-functionalization of virulence functions^8,22^. The maintenance of *AVR_a_* gene variants in the *Bh* population, despite *Mla*-imposed counterselection, suggests that these *Bh* CSEPs have crucial virulence functions. Notably, individual isolates have diversified the *AVR_a_* effector genes to the escape *Mla* recognition and this is predominantly mediated by nucleotide polymorphisms in *AVR_a_* genes but not presence-absence polymorphisms^14–16^, which further underlines this assumption. For example, *Mla1* barley lines are resistant to all *Bh* isolates that carry the *AVR_a1_* effector gene, but not to those isolates that have polymorphisms in the *AVR_a1_* gene that resulted in escape from MLA1 recognition^14,15^. In turn, *AVR_a13_*, whose product is recognized by MLA13, is highly conserved in the *Bh* population and variants with polymorphisms that could mediate the loss of avirulence activity are exceptionally rare^14,15,23^.

We set out to isolate the host targets of *Bh* AVR_A_ effectors and thereby investigate molecular mechanisms employed by agriculturally important biotrophic cereal pathogens to establish host infection. By employing transgenic barley lines stably expressing *AVR_a_* effectors we here identify the host interactome of five AVR_A_ effectors *via* a non-biased *in planta* biotin ligase-based proximity-dependent protein labelling approach. Our data suggests that AVR_A_ RNAse-like effectors interact with divers sets of barley proteins. We identified the barley Strubbelig Receptor Family 3 (HvSRF3) receptor-like kinase as specific interactor of AVR_A13_. AVR_A13_ affects HvSRF3 plasma membrane (PM) abundance and disrupts the association between HvSRF3 and the co-receptor HvBAK1. Importantly, we detect iron-related phenotypes upon AVR_A13_ expression in monocots and dicots, a phenotype that can be genetically linked to SRF3 function. Finally, we show that the AVR_A13_ region required for HvSRF3 interaction overlaps with the residues required for recognition by the MLA13 immune receptor. Together, our results suggest that *Mla* has diversified to specifically detect the residues that underlie RALPH neo-functionalization and intrinsic AVR_A_ virulence functions.

## Results

### Identification of AVR_A_ interactors

To identify the AVR_A_ interactome inside barley cells, we generated stable transgenic barley expressing each of the *Bh AVR_a_* effectors fused to the BirA biotin ligase and a 4myc tag (AVR_A_-BirA). BirA exhibits high specificity in labelling AVR_A_ association partners with biotin that can be isolated using immobilized streptavidin^23^. We obtained barley transformants for *AVR_a1_-BirA*, *AVR_a7_-2-BirA, AVR_a9_-BirA, AVR_a10_-BirA, AVR_a13_-BirA,* and the control line *BirA*. For each sample, barley leaves were infiltrated with 250 µM biotin and proteins were extracted under denaturing conditions prior to streptavidin-mediated isolation of biotinylated proteins and liquid chromatography tandem mass spectrometry (LC-MS/MS)-based identification of isolated proteins (Supplementary Fig. 1a)^24^. Transgene expression and biotinylation of barley proteins by BirA and AVR_A_-BirA were confirmed by anti-myc and Strep-HRP western blots after total protein extraction in triplicate (Supplementary Fig. 1b-1h). This experimental setup was performed independently for each AVR effector and control using two independent transgenic lines. Thereby, we identified 61, 45, 65, 77 and 118 proteins as associators of AVR_A1_, AVR_A7_-2, AVR_A9_, AVR_A10_, and AVR_A13_, respectively (Fig. 1a, Supplementary Fig. 2). We did not observe significant overlaps among interactomes of the different AVR_A_ effector datasets, suggesting that the isolated *Bh* effectors target different sets of proteins in the barley host (Fig. 1b).

**Figure 1:**
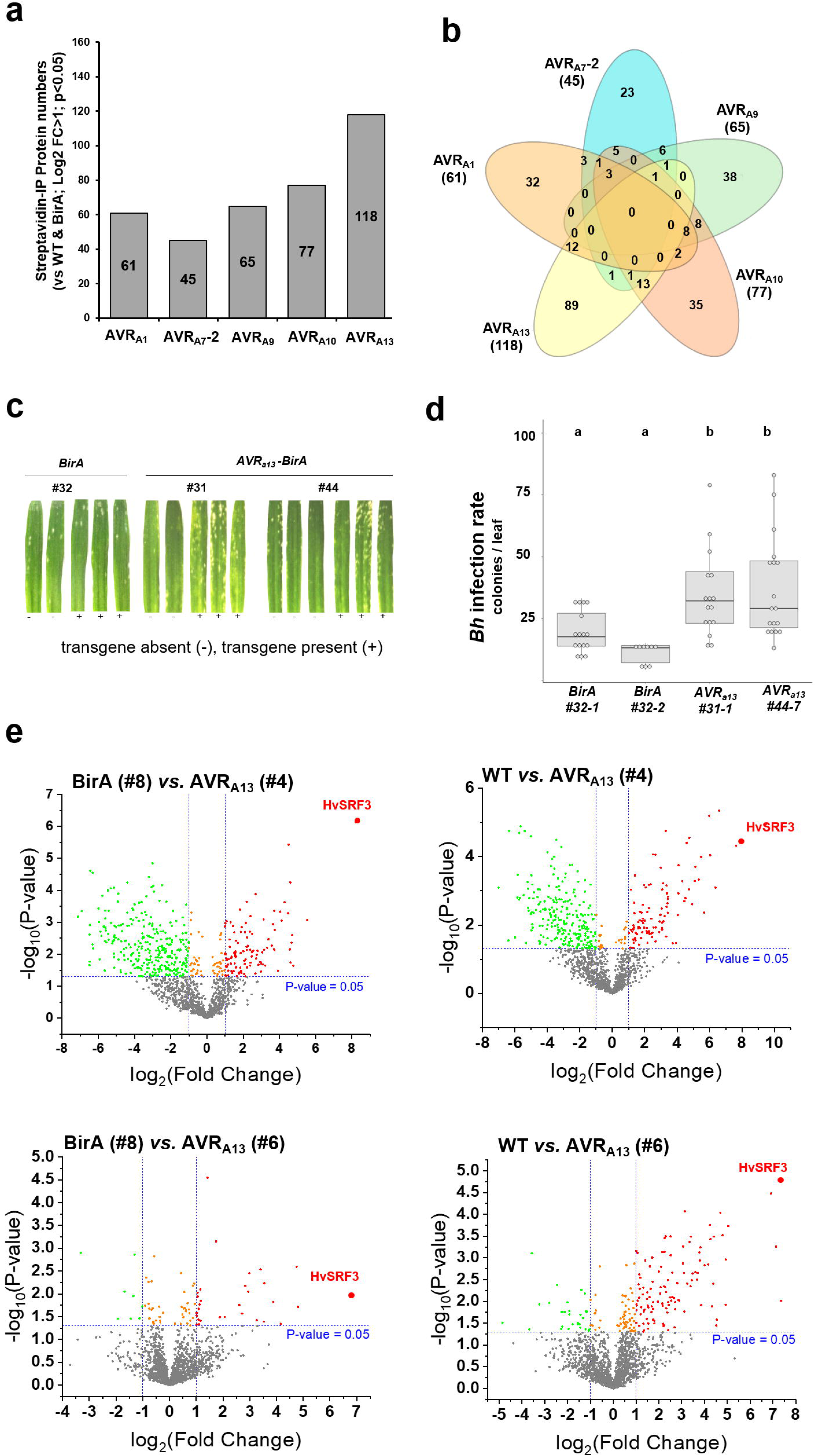
Proximity-dependent protein labelling identifies HvSRF3 as a specific interactor of the *Blumeria hordei* (*Bh*) virulence factor AVR_A13_. **a** Number of barley proteins specifically identified by LC-MS/MS (Log2 FC>1; p<0.05) in proximity labelling samples from transgenic barley expressing AVR_A_ effectors lacking respective signal peptide sequences as indicated and when compared to wild-type (WT) and BirA control samples. **b** Venn diagram investigating the overlap of AVR_A_-interacting proteins from a. c *Blumeria hordei* (*Bh*) disease symptoms on leaves of independent transgenic barley lines (T2) expressing *BirA* or *AVR_a13_-BirA* at five days post infection. Figures shown are representatives of three independent experiments. **d** Quantification of *Bh* colonies at the adaxial and abaxial site of the infected leaves across all replicates of experiments described in **c.** Differences between samples were assessed by Kruskal–Wallis (p=1.177e-05) and subsequent Dunn’s test. Samples marked by identical letters in the plots do not differ significantly (p>0.05) in the Dunn’s test. **e** Volcano plot analysis of proteins specifically identified by LC-MS/MS (at Log2 = 1; p>0.05) in WT (green dots) BirA control (green) and AVR_A13_ (red) proximity labelling samples derived from independent transgenic lines as indicated. Bold dots denote peptides specific to HvSRF3.

### AVR_A13_ associates with barley Strubbelig Receptor Family 3 (SRF3)

*AVR_a13_* is the most highly conserved *AVR_a_* in the global *Bh* population based on the diversification associated with MLA virulence phenotypes^14,15^, suggesting a crucial virulence function for this effector. Accordingly, we consistently detected a significantly higher *Bh* infection rate on barley lines expressing *AVR_a13_* when compared to control plants and respective negative sister lines (Fig. 1c and 1d). This phenotype is likely extrapolated as the level of AVR_A13_ protein may be higher in host cells upon *in planta* expression than upon *Bh* secretion. However, the data demonstrates that *AVR_a13_* expression can indeed exhibit measurable *Bh* virulence activity. We thus set out to further investigate the AVR_A13_ function by mining AVR_A13_-associating proteins. We detected variable enrichment levels of putative AVR_A13_ interactors across proximity labelling experiments using independent *AVR_a13_* transgenic lines (Supplementary Data 1). Yet, across all datasets obtained from both independent transgenic lines, one protein that was identified as barley STRUBBELIG RECEPTOR FAMILY 3-like (HvSRF3) was highly enriched in all the AVR_A13_ samples when compared to both, BirA and wild-type control samples (Fig. 1e, Supplementary Data 1).

SRF3 is a PM-localized leucine-rich repeat receptor-like kinase (LRR-RLK) with a unique N-terminal STRUBBELIG (SUB) domain^25^. Four barley LRR-RLKs are annotated as HvSRF3-like^26^, and we named these HvSRF3-Like1 (HvSRF3-L1, LOC123409827), HvSRF3-Like2 (HvSRF3-L2, LOC123401201), HvSRF3-Like3 (HvSRF3-L3, LOC123425675) and HvSRF3-Like4 (HvSRF3-L4, LOC123403104) (Fig. 2, Supplementary Fig. 3, Supplementary Fig. 4). By searching available barley databases (NCBI, EnsemblPlants and Uniprot), we identified a total of 12 *HvSRF*-like genes. Sequence comparison suggested that these are orthologous to *A. thaliana* SRF2, SRF4, SRF5, SRF6, SRF7 and SRF8 and that the previously annotated HvSRF3 proteins are homologous to *A. thaliana* SRF3, although we also detected protein sequence homology between HvSRF3-L4 and AtSRF9 (AtSUB, Fig. 2a and 2b, Supplementary Fig. 3)^25^.

**Figure 2:**
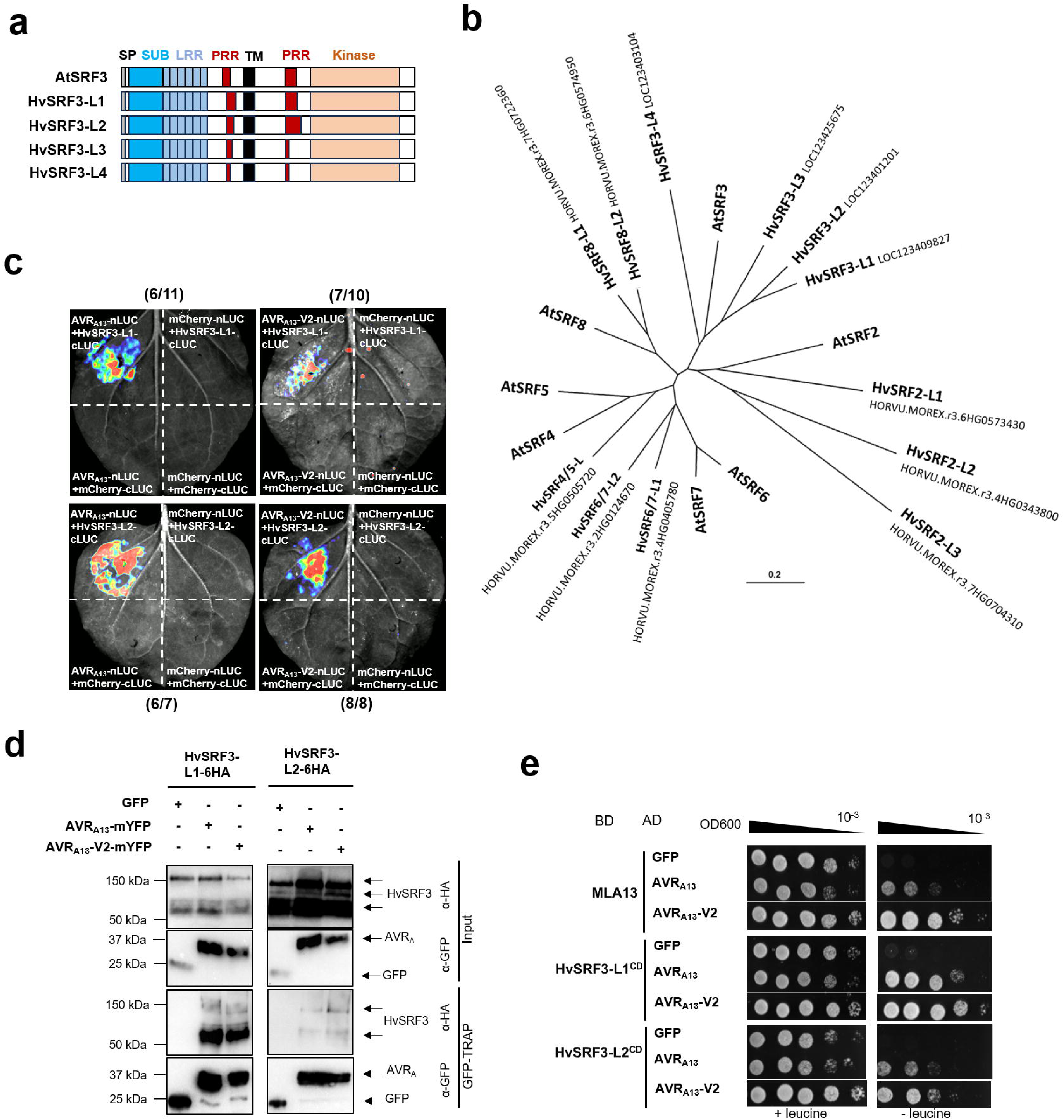
*Blumeria hordei* (*Bh*) AVR_A13_ can interact with the cytoplasmic domain (CD) of HvSRF3. **a** Domain structure of *Arabidopsis thaliana* and barley Strubbelig Receptor Family 3 (SRF3). **b** Phylogeny (Neighbor-Joining tree) of *A. thaliana* and barley Strubbelig Receptor Family members (HvSRFs). **c and d** *Nicotiana benthamiana* leaves were transiently transformed with the constructs indicated. At three days post transformation, protein-protein interaction was determined by the complementation of the luciferase (LUC) reporter by nLUC and cLUC-fused protein upon addition of substrate (c) and GFP-TRAP-based immunoprecipitation of GFP or mYFP tagged proteins followed by anti-GFP and anti-HA western blot analysis (d). Figures shown are representatives of at least three independent experiments. **e** Yeast cells were transformed with B42 activation domain (AD)- and HA tag-fused constructs encoding GFP or AVR_A13_ variants together with LexA binding domain (BD)-fused constructs encoding MLA13, HvSRF3-L1 cytoplasmic domain (HvSRF3-L1^CD^) or HvSRF3-L2 cytoplasmic domain (HvSRF3-L2^CD^). Growth of transformants was determined on selective growth medium with leucine (+leucine), and interaction of proteins was determined by leucine reporter activity reflected by growth of yeast on selective medium lacking leucine (-leucine). Figures shown are representatives of at least three independent transformation experiments and pictures were taken three days after drop out. Signal peptides were depleted from all effector-encoding constructs.

In the AVR_A13_ proximity labelling datasets, we only identified peptides specific to HvSRF3-L1 and HvSRF3-L2 (Supplementary Data 1). To determine if HvSRF3-L1 and HvSRF3-L2 indeed interact with AVR_A13_, we tested their association with AVR_A13_ and its rare naturally occurring virulent variant AVR_A13_-V2^14,15,23^ by luciferase complementation (split-luciferase assay) upon co-expression of respective constructs in *Nicotiana benthamiana* leaves. We detected luciferase activity upon co-expression of *AVR_a13_* and the virulent variant *AVR_a13_-V2* with both *HvSRF3-L1* and *HvSRF3-L2* (Fig. 2c). Additionally, immunoprecipitation of AVR_A13_-mYFP and AVR_A13_-V2-mYFP but not GFP co-isolated HvSRF3-L1-6HA and HvSRF3-L2-6HA (Fig. 2d). Both HvSRF3 proteins appear n-terminally fragmented, and HvSRF3-L1-6HA protein is less stable than HvSRF3-L2 upon *in planta* expression (Fig. 2d; Supplementary Fig. 5a). As an additional control, we employed the common co-receptor BAK1 from barley (HvBAK1, EF216861.1^27^), which was not detected in the proximity-dependent protein labelling dataset. Consistent with this, we did not detect a steady interaction of AVR_A13_ with HvBAK1 (Supplementary Figs. 5b and 5c).

Because AVR_A13_ is recognized by MLA13 inside barley cells^14,15^, it is plausible to assume that AVR_A13_ interacts with the cytoplasmic domain (CD) of HvSRF3. To test this, we performed yeast two-hybrid (Y2H) assays with AVR_A13_ and the cytoplasmic domains of HvSRF3-L1 (HvSRF3-L1^CD^) and HvSRF3-L2 (HvSRF3-L2^CD^). We used MLA13 as a control and as expected, both AVR_A13_ and AVR_A13_-V2, but not GFP interacted with MLA13 (Fig. 2e)^14,23^. We also detected interaction of AVR_A13_ and AVR_A13_-V2 with HvSRF3-L1^CD^ and HvSRF3-L2^CD^ (Fig. 2e, Supplementary Fig. 5d).

### Effector binding of HvSRF3 is specific to AVR_A13_ and mediated by the HvSRF3 kinase domain

*Bh* AVR_A_ effectors other than AVR_A13_ did not biotinylate any of the HvSRF3-Like proteins in the proximity labelling experiment (Fig. 1b, Supplementary Data 1). To determine if indeed only AVR_A13_ can bind HvSRF3, we also tested if other AVR_A_ effectors can associate with HvSRF3 in directed protein-protein interaction assays. In agreement with the proximity labelling experiments, we did not detect interaction of HvSRF3-L1 or HvSRF3-L2 with any of the other AVR_A_ effectors, despite stable detection of all proteins (Fig. 3a, Supplementary Fig. 6a). We conclude that HvSRF3-L1 and HvSRF3-L2 are specific targets of AVR_A13_.

**Figure 3:**
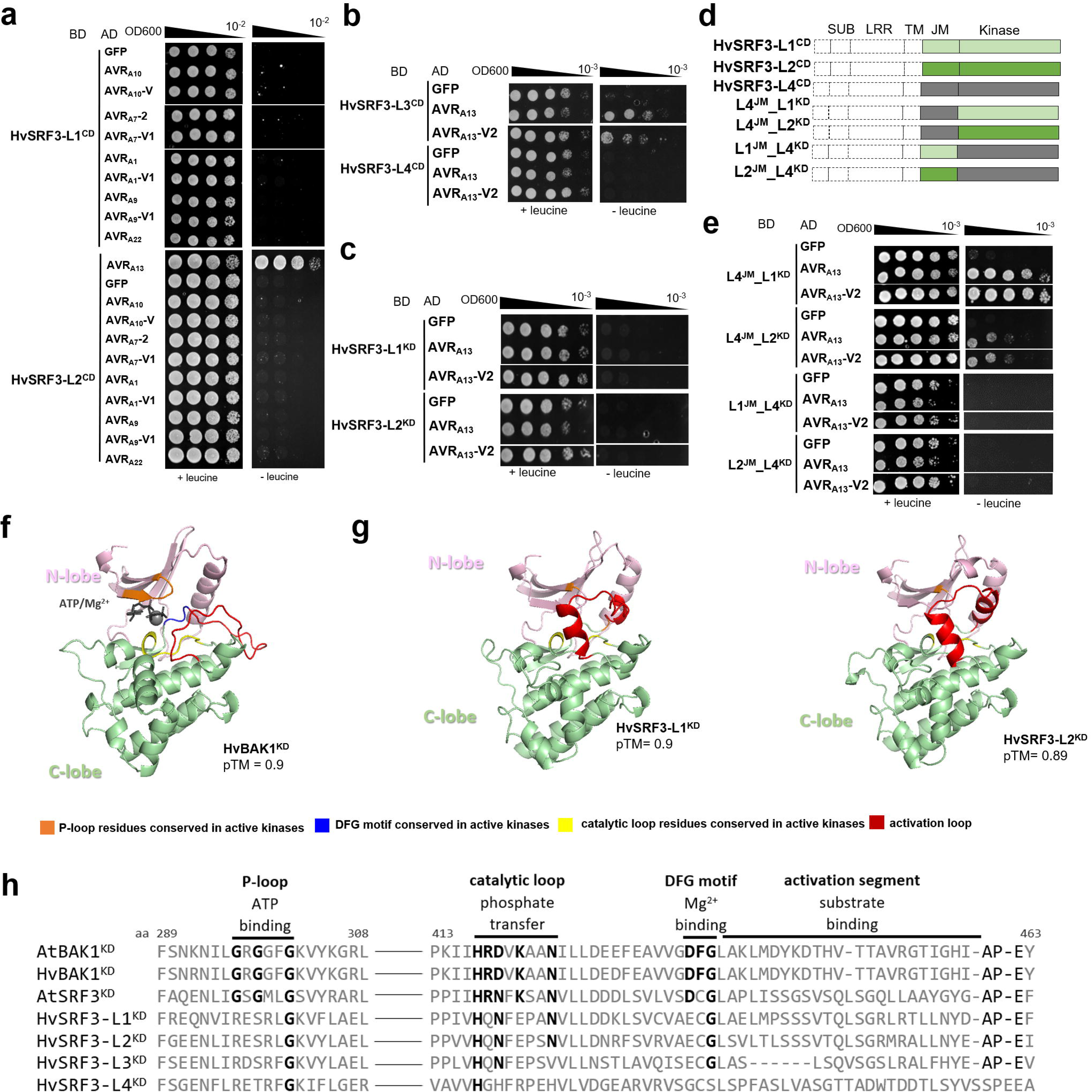
The HvSRF3 association is specific to AVR_A13_ and facilitated by the HvSRF3 kinase domain. **a to c** Yeast cells were transformed with B42 activation domain (AD)- and HA tag-fused constructs encoding GFP or AVR_A_ variants together with LexA binding domain (BD)-fused constructs encoding either of the HvSRF3 cytoplasmic domains (HvSRF3^CD^) (a, b) or HvSRF3 kinase domains (HvSRF3^KD^) (c) as indicated. Growth of transformants was determined on selective growth medium with leucine (+leucine), and interaction of proteins was determined by leucine reporter activity reflected by growth of yeast on selective medium lacking leucine (-leucine). **d** Domain architecture of HvSRF3 isoforms and chimeric HvSRF3 constructs composed of the HvSRF3 isoform L1 (pale green), L2 (green) or L4 (grey) juxtamembrane domain (JM) and kinase domain (KD). **e** Yeast cells were transformed with B42 activation domain (AD)- and HA tag-fused constructs encoding GFP or AVR_A13_ variants together with LexA binding domain (BD)-fused constructs encoding chimeric HvSRF3 constructs from c. Protein interaction was determined as described in a. Signal peptides were depleted from all effector-encoding constructs. Figures shown are representatives of at least three independent transformation experiments and pictures were taken three days after drop out. **f and g** AlphaFold models of ATP and Mg^2+^-bound HvBAK1^KD^ (pTM =0.9) (f), HvSRF3-L1^KD^ (pTM =0.9) and HvSRF3-L2^KD^ (pTM =0.89) (g). The N-lobe, C-lobe, P-loop, DFG motif, activation loop, catalytic loop in HvBAK1^KD^ or respective residues in HvSRF3-L1^KD^/HvSRF3-L2^KD^ are coloured cyan, pale green, orange, blue, red and yellow, respectively. **h** Sequence alignment of critical kinase motifs in AtBAK1^KD^, HvBAK1^KD^, AtSRF3^KD,^ HvSRF3-L1^KD^, HvSRF3-L2^KD^, HvSRF3-L3^KD^, HvSRF3-L4^KD^. Residues involved in phosphotransferase activity are highlighted in bold.

We did also not identify HvSRF3-L3 or HvSRF3-L4 in the AVR_A13_ proximity labelling samples (Supplementary Data 1). To determine the general capacity of AVR_A13_ to bind HvSRF3-L3 and HvSRF3-L4, we performed Y2H assays with *HvSRF3-L3^CD^* and *HvSRF3-L4^CD^*and found that HvSRF3-L4^CD^ could not interact with AVR_A13_ and AVR_A13_-V2 (Fig. 3b, Supplementary Fig. 6b). In turn, we detected association of AVR_A13_ and AVR_A13_-V2 with HvSRF3-L3^CD^ but this interaction appeared weaker than the association with HvSRF3-L1^CD^ and HvSRF3-L2^CD^ (Fig. 3b, Supplementary Fig. 6c). HvSRF3-L4 expression is generally not detectable, and unlike HvSRF3-L1 and HvSRF3-L2, HvSRF3-L3 is not upregulated upon pathogen attack (http://barleyexp.com/common.html), suggesting that HvSRF3-L3 and HvSRF3-L1/HvSRF3-L2 functions do not overlap. Thus, we speculate that *Bh* employs AVR_A13_ to primarily target HvSRF3-L1^CD^ and HvSRF3L2^CD^ but note that AVR_A13_ can in principle bind HvSRF3 upon overexpression of both proteins.

The cytoplasmic regions of HvSRF3-like proteins consist of the C-terminal kinase domains (KDs) and the juxtamembrane domains (JMs) that encompass a promiscuous proline-rich-region (PRR) (Fig. 2, Supplementary Fig. 3, Supplementary Fig. 4)^25^. Sequence analysis demonstrates high variability within the JM domain across the HvSRF3-Like proteins with HvSRF3-L4^JM^ being shorter than the JMs of the other three HvSRF3-Like proteins (Supplementary Fig. 4). To understand if the differences in the JM domain account for the complete lack of interaction between AVR_A13_ and HvSRF3-L4, we tested if AVR_A13_ can bind the HvSRF3-L1^KD^ and HvSRF3-L2^KD^, which represent the CDs without respective JMs. Neither AVR_A13_ nor AVR_A13_-V2 showed interaction with HvSRF3-L1^KD^ or L2^KD^ (Fig. 3c, Supplementary Fig. 6d). To further clarify the roles of the individual HvSRF3 domains in mediating the AVR_A13_ association, we generated hybrid constructs exchanging the JM domains of HvSRF3-L1 and HvSRF3-L2 with the HvSRF3-L4^JM^ and *vice versa* (Fig. 3d). Only the hybrids encoding the HvSRF3-L1^KD^ or HvSRF3-L2^KD^ interacted with AVR_A13_ (Fig. 3d and 3e, Supplementary Fig. 6e). We conclude that AVR_A13_ requires the JM of HvSRF3 for interaction in yeast, but that specificity of the interaction between AVR_A13_ with HvSRF3-L1 and HvSRF3-L2 is mediated by the respective HvSRF3^KD^s. Interestingly, kinase activity could not be detected for SRF3 and other SRFs from *A. thaliana*^28–31^. We therefore evaluated the general propensity of the HvSRF3^KD^s for putative phosphorylation capacity by assessing the conservation of key kinase residues. We also visualized the KD structures (AlphaFold3) of HvSRF3 and the kinase active receptor HvBAK1 for comparison (Fig. 3f to 3h). The analyses demonstrate that in addition to the protein polymorphisms within the conserved catalytic loop of AtSRF3 when compared to the kinase active BAK1^KD^s, HvSRF3^KD^s lack both, the GxGxxG motif of the P-loop and the DFG motif. Correspondingly, neither Mg^2+^ nor ATP is modelled into the HvSRF3 binding pockets (Fig. 3f to 3h, Supplementary Fig. 6f)^32^. We conclude that, similar to AtSRF3 and AtSRF9, the KDs of HvSRF3s are most likely pseudokinases. SRF3 and other SRFs may thus act as molecular scaffolds for other RLKs or rely on the kinase activity of associating co-receptors with active kinases and we speculate that either of these putative HvSRF3 functions is manipulated by AVR_A13_.

### AVR_A13_ affects HvSRF3-HvBAK1 interaction and HvSRF3 arrangement at the plasma membrane

Cell surface-localized LRR-RLKs frequently rely on the co-receptor BAK1/SERK3 (SOMATIC EMBRYOGENESIS RECEPTOR KINASE3) or other SERK family members to transduce signals across the PM^33,34^. Because multiple SRF members have been reported to associate with BAK1^35–37^, we hypothesized that HvSRF3 might rely on HvBAK1 or related co-receptors to transduce unknown signals across the PM. We therefore determined if HvSRF3 and HvBAK1 associate upon co-expression. For this, we primarily focused on HvSRF3-L2 (hereafter referred to as HvSRF3), because the HvSRF3-L1 protein is poorly detectable upon transient *in planta* expression (Fig. 2d; Supplementary Fig. 5a). Y2H analysis involving HvBAK1^CD^ and HvSRF3-L2^CD^ (Fig. 4a, Supplementary Fig. 7a) and split-luciferase assays of full length HvBAK1 and HvSRF3-L2 in *N. benthamiana* leaves confirmed interaction of HvSRF3 and HvBAK1 (Fig. 4b). In agreement, we detected fluorescence signals *via* YFP complementation by co-expression of HvBAK1 fused to the C-terminal domain of YFP (HvBAK1-YFP^C^) and HvSRF3 fused to the N-terminal domain of YFP (HvSRF3-YFP^N^, Fig. 4c).

**Figure 4:**
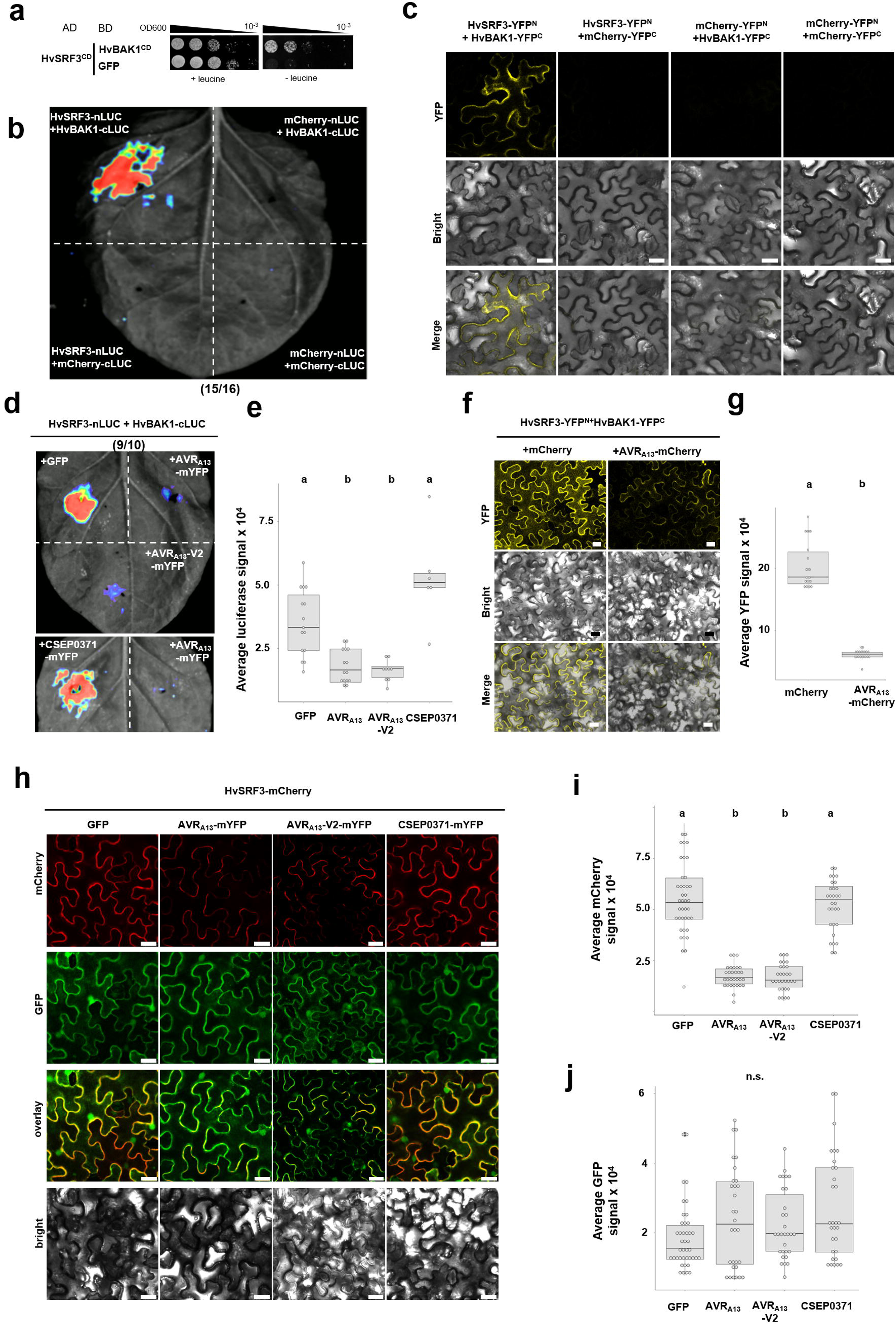
AVR_A13_ affects HvSRF3 protein localization and its interaction with HvBAK1. **a** Yeast cells were transformed with B42 activation domain (AD)- and HA tag-fused constructs encoding the HvSRF3^CD^ together with the LexA binding domain (BD)-fused construct encoding the cytoplasmic domain of HvBAK1 (HvBAK1^CD^). Growth on selective medium with leucine (+leucine) confirmed transformation, while growth on medium lacking leucine (-leucine) indicated protein–protein interaction *via* activation of the leucine reporter. Photographs were taken three days after drop-out. **b** *Nicotiana benthamiana* leaves were transiently co-transformed with C-terminally nLUC-fused and C-terminally cLUC-fused constructs as indicated. At three days post transformation, protein-protein interaction was determined by the complementation of the luciferase (LUC) reporter by chemiluminescence upon addition of substrate. Pictures shown are representative results from at least three independent biological replicates. **c** *N. benthamiana* leaves were transiently co-transformed with C-terminally YFP^N^-fused and C-terminally YFP^C^-fused constructs as indicated. At two days post-transformation, protein-protein interaction was determined by the complementation of the YFP fluorescence signals using a Leica TCS SP8 confocal laser scanning microscope. Pictures shown are representative results from at least three independent biological replicates. Scale bars: 25 µm. **d** *N. benthamiana* leaves were transiently co-transformed with constructs encoding GFP or mYFP tagged effector constructs as indicated together with C-terminally nLUC-fused HvSRF3 and C-terminally cLUC-fused HvBAK1 constructs. At three days post transformation, protein-protein interaction was determined by the complementation of the luciferase (LUC) reporter by chemiluminescence upon addition of substrate. Pictures shown are representative results from at least three independent biological replicates. **e** Quantification of luciferase signals across all replicates of experiments described in d. Differences between samples assessed by Kruskal-Wallis (p=1.324e-05) and subsequent Dunn’s test. Samples marked by identical letters in the plots do not differ significantly (p>0.05) in the Dunn’s test. **f** *N. benthamiana* leaves were transiently transformed with constructs encoding mCherry or AVR_A13_-mCherry together with C-terminally YFP^N^-fused HvSRF3 and C-terminally YFP^C^-fused HvBAK1constructs as indicated. At two days post transformation, protein-protein interaction was determined by complementation of the YFP fluorescence signal using a Leica TCS SP8 confocal laser scanning microscope. Pictures shown are representative results from at least three independent biological replicates. Scale bars: 25 µm. **g** Quantification of luciferase signals across all replicates of experiments described in f. Differences between samples assessed by Kruskal–Wallis (p=3.015e-08) and subsequent Dunn’s test. Samples marked by identical letters in the plots do not differ significantly (p>0.001) in the Dunn’s test. **h** *N. benthamiana* leaves were transiently transformed with constructs encoding HvSRF3-mCherry together with constructs encoding GFP, AVR_A13_-mYFP, AVR_A13_-V2-mYFP or CSEP0371-mYFP. GFP/YFP and mCherry fluorescence signals were determined using a Leica TCS SP8 confocal laser scanning microscope two days post-transformation. Pictures shown are representative results from at least three independent biological replicates. Scale bars: 25 µm. **i** Quantification of mCherry signals across all replicates of experiments described in h. **j** Quantification of GFP/YFP signals across all replicates of experiments described in h. Differences between samples were assessed by Kruskal–Wallis (p= 2.2e-16 for I and p= 0.09609 for J) and subsequent Dunn’s test. Samples marked by identical letters in the plots do not differ significantly (p>0.001) in the Dunn’s test. n.s. = no significant difference. Signal peptides were depleted from all effector-encoding constructs throughout.

Several pathogen effectors manipulate receptor signaling outputs by modulating association with the BAK1 co-receptor^38^. We therefore investigated if AVR_A13_ can manipulate association of HvSRF3 with HvBAK1. In split-luciferase assays, signals mediated by HvBAK1-cLUC and HvSRF3-nLUC were abrogated in the presence of AVR_A13_-mYFP and AVR_A13_-V2-mYFP but not GFP (Fig. 4d and 4e, Supplementary Fig. 7b). We also tested the specificity of this effect by employing CSEP0371, which is the *Bh* effector most closely related to AVR_A13_ (also known as CSEP0372)^8^ and found that CSEP0371-mYFP does not affect luciferase signal of HvBAK1-cLUC and HvSRF3-nLUC (Fig. 4d and 4e, Supplementary Fig. 7b). Similar results were obtained in a YFP complementation assay, where the YFP signal that was detectable upon co-expression of HvBAK1-YFP^C^ and HvSRF3-YFP^N^ was largely diminished in the presence of AVR_A13_-mCherry but not mCherry alone (Fig. 4f and 4g, Supplementary Fig. 7c and 7d). We detected a fluorescent signal at the PM but also in the cytosol and also concentration of the receptor complex in specific regions at the PM in the presence of AVR_A13_ (Fig. 4f and 4g, Supplementary Fig. 7e). Thus, we tested if AVR_A13_ is able to mediate changes to HvSRF3 PM abundance, a process associated with AtSRF3 function in *A. thaliana* roots^39^. We co-expressed *HvSRF3-mCherry* with either effector or the *GFP* control. HvSRF3 PM signals but not total HvSRF3 protein levels are reduced in the presence of AVR_A13_ and AVR_A13_-V2 but not CSEP0371 when compared to the GFP control (Fig. 4h to 4j, Supplementary Fig. 7f). Effector signals were also detected in the nucleus (Fig. 4h, Supplementary Fig. 7c and 7e). We cannot clarify if AVR_A13_-medited changes to HvSRF3 PM organization is the cause or a consequence of abrogated HvSRF3-HvBAK1 interaction. Yet, upon transient co-expression of *HvSRF3* and *HvBAK1*, it appears that HvSRF3 does not significantly affect HvBAK1 protein or HvBAK1-mCherry signals at the plasma membrane (Supplementary Fig. 7g and 7h). We conclude that the AVR_A13_ -HvSRF3 interaction is associated with changes in HvSRF3 PM localization and affects the ability of HvSRF3 to interact with HvBAK1.

### AVR_A13_-mediated modulation of SRF3 is conserved between monocots and dicots

Despite high levels of conservation across land plants^40^, the molecular functions of SRF members are not fully understood. Recently, rice SRF3 has been implicated in regulating chitin-induced PTI responses^37^. In protoplasts derived from rice lines overexpressing OsSRF3, chitin-induced production of reactive oxygen species (ROS) was elevated, and this was affected by the effector UgsL from the fungal pathogen *Ustilaginoidea virens*^37^. To determine if AVR_a13_ modulates typical PTI output to mediate pathogen virulence, we assayed PAMP-mediated ROS production using the stable *AVR_a13_*-expressing barley lines with increased susceptibility to *Bh*. We did not detect any consistent differences in chitin- or flg22-mediated ROS production between control or *AVR_a13_*-expressing barley lines (Supplementary Fig. 8).

In *A. thaliana*, AtSRF3 regulates iron homeostasis and nutritional immunity^29,39^. AtSRF3-mediated sensing of low iron signals is associated with reduced AtSRF3 levels at the PM. In addition, low iron-mediated root growth inhibition is abolished in *A. thaliana srf3* lines ^39^. We therefore decided to genetically determine if *AVR_a13_*-expression can affect SRF3 phenotypes under low iron in *A. thaliana*. We found that AVR_A13_ can indeed bind AtSRF3 using split-luciferase, co-immunoprecipitation, and Y2H assays (Fig. 5a to 5c Supplementary Fig. 9a and 9b). Thus, we generated homozygous *AVR_a13_*-expressing *A. thaliana* Col-0 lines (Fig. 5d, Supplementary Fig. 9c) and determined low iron-mediated root growth response^39^. At physiological iron levels, *AVR_a13_*expressing *A. thaliana* seedlings did not show root growth inhibition (Fig. 5e). However, the low iron-induced root growth inhibition response was impaired in *AVR_a13_*-expressing lines, and this mimics *srf3* mutant phenotypes (Fig. 5f)^39^. Together, the data suggest that AVR_A13_ can impair SRF3 function in regulating iron deficiency responses in dicots.

**Figure 5:**
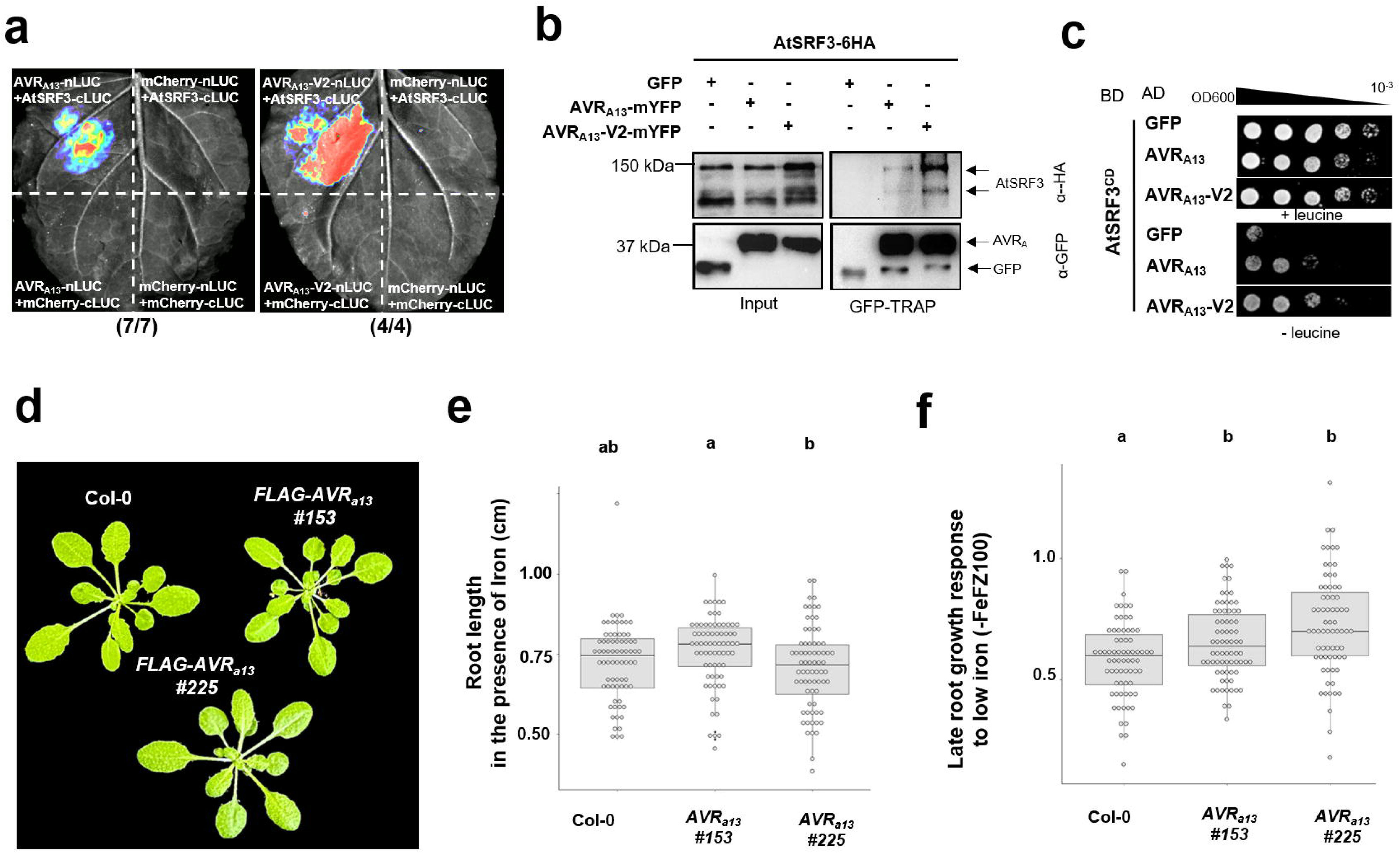
AVR_A13_ can associate with *Arabidopsis thaliana* SRF3 (AtSRF3) and modulates plant responses to differential iron levels in *A. thaliana*. **a and b** *Nicotiana benthamiana* leaves were transiently transformed with the constructs indicated. At three days post transformation, protein-protein interaction was determined by the complementation of the luciferase (LUC) reporter by chemiluminescence upon addition of substrate (a) and GFP-TRAP-based immunoprecipitation of GFP or mYFP tagged proteins followed by anti-GFP and anti-HA western blot analysis (b). Figures shown are representatives of at least three independent experiments. **c** Yeast cells were transformed with B42 activation domain (AD)- and HA tag-fused constructs encoding GFP or AVR_A13_ variants together with LexA binding domain (BD)-fused constructs encoding the AtSRF3 cytoplasmic domain (AtSRF3^CD^). Growth of transformants was determined on selective growth medium with leucine (+leucine), and interaction of proteins was determined by leucine reporter activity reflected by growth of yeast on selective medium lacking leucine (-leucine). Figures shown are representatives of at least three independent transformation experiments and pictures were taken three days after drop out. d *A. thaliana* rosettes of Col-0 and *AVR_a13_*-expressing lines grown under standard conditions. **e** Root length of *A. thaliana* lines from d at eight days grown *in vitro* under iron sufficient growth conditions. **f** Late root growth response (Platre *et al.,* 2024) of Col-0 wild-type and *AVR_a13_*-expressing *A. thaliana* lines grown for four days with (50 µM FeNaEDTA) or without iron (-FeFZ100). Differences between lines were assessed by analysis of variance (ANOVA p=1.75E-05) and subsequent Tukey post hoc test. Samples marked by identical letters in the plots do not differ significantly (p>0.05) in the Tukey test. Values were obtained from three independent experiments. Signal peptides were depleted from all effector-encoding constructs throughout.

Iron uptake and transport strategies, as well as iron deficiency responses differ greatly between dicots such as *A. thaliana* and monocots such as barley. Compared with roots, our understanding of iron transport and homeostasis in leaf tissues is limited^42–44^. Powdery mildew attack has indeed been shown to cause local intracellular iron depletion and extracellular increase of ferric iron, which is associated with the production of reactive oxygen species such as H_2_O_2_ and nitric oxide^41^. However, it is unclear how fungal penetration-induced alterations to host iron homeostasis affect the outcome of fungal penetration attempts. This and elevated *AVR_a13_* expression post penetration^8,9^ suggests *AVR_a13_*-mediated modulation of host processes upon successful penetration. We therefore decided to determine how differential iron levels actually affect powdery mildew disease development beyond fungal penetration. For this, we infected barley grown at differential iron levels with *Bh*. Plants grown under low iron conditions (10 µM Fe-EDTA, which represents 10 x less iron than the control sample) were not significantly impacted in *Bh* susceptibility, whereas barley grown at high levels of iron (250 µM FeEDTA, which represents 2.5 x more iron than the control sample) showed reduced *Bh* infection rates (Supplementary Fig. 10a). To mimic a putative host-induced iron accumulation at the site of infection and counteract intracellular iron depletion without disturbing the whole plant’s iron status, we applied chelated iron (FeEDDHA) to barley leaves prior to *Bh* infection. We found that this also leads to reduced *Bh* disease symptoms (Fig. 6a and 6b). Importantly, in contrast to controls, barley lines expressing *AVR_a13_* did not exhibit FeEDDHA-induced reduction of fungal proliferation (Fig. 6c and 6d), whereas barley lines expressing *AVR_a9_* were sensitive to FeEDDHA application, similar to control lines (Supplementary Fig. 10b and 10c). From this, we conclude that high levels of iron negatively influence *Bh* proliferation and that AVR_A13_ but not AVR_A9_, which cannot bind HvSRF3 (Fig. 1b, Fig. 3a), desensitizes fungal susceptibility to high iron levels. To further determine if this AVR_A13_-specific function can indeed be linked to its interaction with HvSRF3, we used a CRISPR-Cas9 approach to generate plants mutated in *HvSRF3* in the genetic background cv. Golden Promise (wild-type). We obtained two independent barley *srf3* lines with double-homozygous deletions in *HvSRF3-L1* and *HvSRF3-L2*. The respective lines have a single base pair and a double base pair deletion, respectively, leading to early stop codons in both genes (hereafter named *srf3* lines, Supplementary Fig. 10d) and did not display consistent growth phenotypes under standard growth conditions (Supplementary Fig. 10e). To survey *Bh* colonization, we inoculated the control and both *srf3* mutant lines and tested the effect of FeEDDHA addition. In the absence of FeEDDHA, we consistently detected a significantly higher fungal infection rate on the *srf3* lines when compared to control lines and importantly, the *srf3* barley lines were insensitive to the decreased fungal infection upon FeEDDHA application, similar to *AVR_a13_*-expressing lines (Fig. 6e and 6f). Notably, similar to *AVR_a13_*-expressing barley lines, neither chitin-nor flg22-induced ROS production was affected in the barley *srf3* lines (Supplementary Fig. 10f to 10i). We conclude that HvSRF3 negatively regulates *Bh* infection and that *AVR_a13_* lines mimic *srf3* lines in their inability to mediated iron-induced *Bh* restriction. We therefore conclude that, for promoting virulence, AVR_A13_ negatively regulates HvSRF3 and hypothesize that AVR_A13_ targets HvSRF3 to manipulate iron-associated processes that are independent from chitin-induced ROS production.

**Figure 6:**
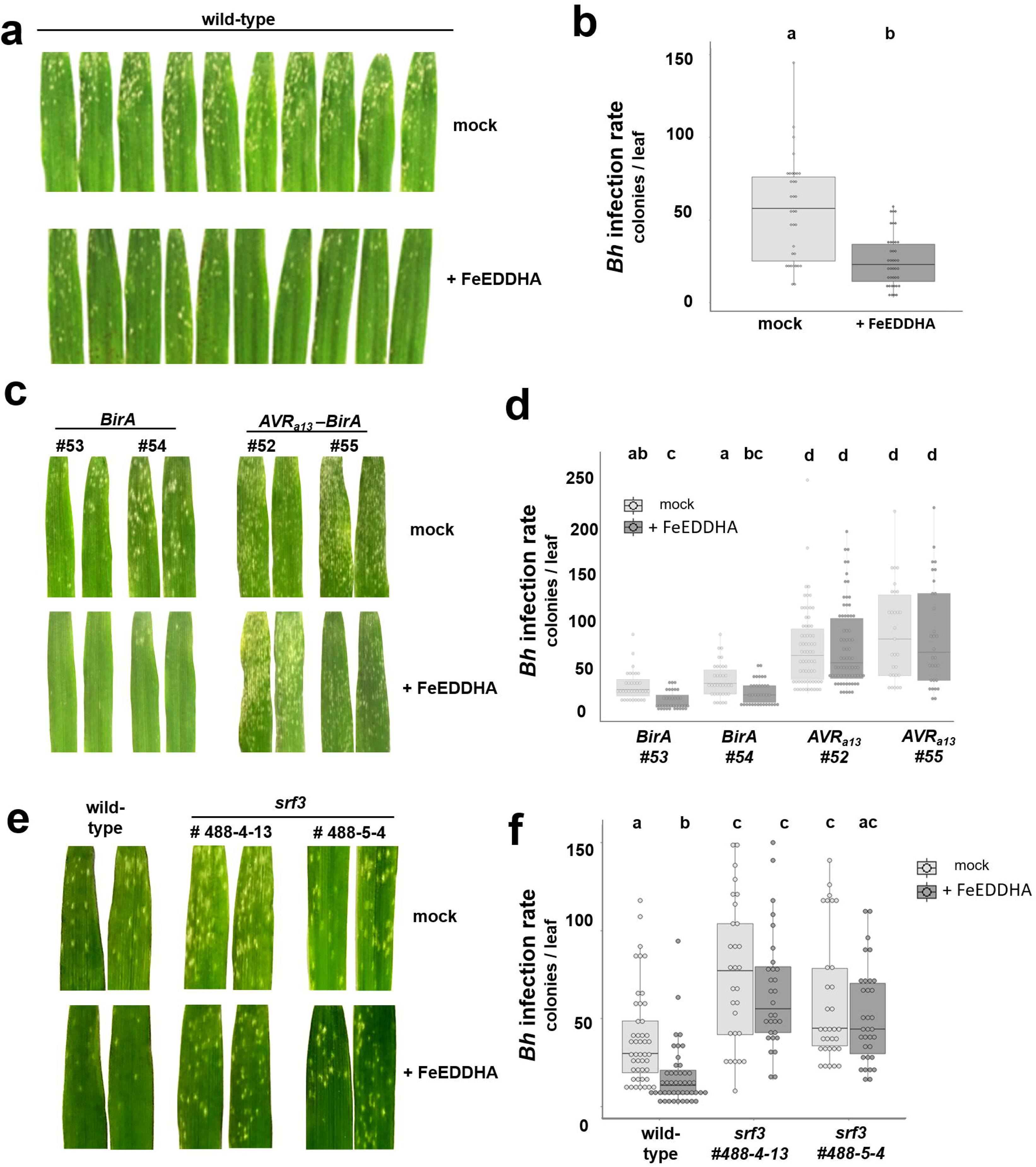
AVR_A13_ and HvSRF3 modulate high iron sensitivity of *Bh* infection of barley. **a and b** Leaves of barley cultivar Golden Promise Fast plants were pre-treated with 0.5 g/L FeEDDHA or water (mock) prior to infection with *Blumeria hordei* (*Bh*). Fungal growth was determined at five days post-infection. Figures shown are representatives of at least three independent experiments (a) and quantification of *Bh* colonies was performed from pictures takes of the adaxial and abaxial site of the infected leaves across all replicates of experiments described. Differences between treatments were assessed by the Kruskal–Wallis test (p=1.352e-05) (b). **c and d** *Bh* disease symptoms on leaves of independent transgenic barley lines (T3) expressing *BirA* or *AVR_a13_-BirA* at five days post-infection upon treatment with Tween (mock) or 0.5 g/L FeEDDHA. Figures shown are representatives of at least three independent experiments (c). Quantification of *Bh* colonies (d) was performed from pictures takes of the adaxial and abaxial site of the infected leaves across all replicates of experiments described in c. Differences between treatments were assessed by the Kruskal–Wallis test (p<2.2e-16) and subsequent Dunn’s test. Samples marked by identical letters in the plots do not differ significantly (p>0.05) in the Dunn’s test. **e and f** *Bh* disease symptoms on leaves of independent homozygous *srf3* barley CRISPR mutant lines at five days post infection upon treatment with water (mock) or 0.5 g/L FeEDDHA. Figures shown are representatives of at least three independent experiments (e). Quantification of *Bh* colonies was performed from pictures takes of the adaxial and abaxial site of the infected leaves across all replicates of experiments described in e. Differences between treatments were assessed by Kruskal–Wallis test (p<2.2e-16) and subsequent Dunn’s test. Samples marked by identical letters in the plots do not differ significantly (p>0.05) in the Dunn’s test. Signal peptides were depleted from all effector-encoding constructs throughout.

### AVR_A13_ residues required for SRF3 interaction are recognized by MLA13

All AVR_A_ effectors analyzed here belong to the *Bh* RALPH effector family, for which it was postulated that multiple members can target the same host processes to promote virulence^18,45^. AVR_A13_ (also known as CSEP0372) has three family members in *Bh*, namely CSEP0371, CSEP0373 and CSEP0374 (Fig. 7a and 7b)^8^. CSEP0371 and CSEP0374 are expressed during *Bh* infection of barley, but CSEP0373 is not^9^. We already demonstrated that unlike AVR_A13_, CSEP0371 does not affect HvSRF3 phenotypes (Fig. 4). To directly determine CSEP family-wide conservation of *Bh* effector targets, we tested interaction of CSEP0371 and CSEP0374 with the HvSRF3-L1^CD^, which showed the strongest interaction with AVR_A13_ in Y2H assays (Fig. 2e). We failed to detect interaction of CSEP0371 and CSEP0374 with HvSRF3-L1^CD^ in Y2H (Fig. 7c, Supplementary Fig. 11a). We then made use of this to identify the HvSRF3-binding residues in AVR_A13_. For this, we generated eight chimeric AVR_A13_ / CSEP0371 constructs (chimera 1-8, Fig. 6d). Interaction of chimera 6 and chimera 8 but not chimera 4 with the HvSRF3^CD^ in the Y2H assay suggests that a central loop region of the AVR_A13_ protein mediates HvSRF3 interaction (Fig. 7e, Supplementary Fig. 11b). Interestingly, this region contains seven of the 11 residues previously identified as MLA13 contact residue^46^. Based on these observations, we hypothesized that these MLA13 contact residues may also play a role in the interaction of AVR_A13_ with HvSRF3. We therefore exchanged corresponding AVR_A13_ residues to CSEP0371 residues (chimera 10) and *vice versa* (chimera 9). Additionally, we generated chimeric constructs to more specifically define the region of AVR_A13_ that mediates HvSRF3 interaction (chimera 11, 12 and 13). As determined by the use of chimera 9, the introduction of the AVR_A13_-MLA13 contact residues into CSEP0371 alone was insufficient for transferring the ability of AVR_A13_ to bind HvSRF3. On the other hand, the elimination of the MLA13 contact residues in AVR_A13_ in chimera 10 disrupted effector binding to HvSRF3 (Fig. 7e, Supplementary Fig. 11b), implying that the MLA13 contact residues of AVR_A13_ play a crucial role for association with HvSRF3. Chimera 10 to chimera 13 narrowed down the functional region for HvSRF3 interaction to AVR_A13_ amino acid (aa) E44 to aa N70, which is largely composed of a central loop region (Fig. 7i and 7j). Further exchanges of individual AVR_A13_ amino acids to corresponding CSEP0371 residues in this region (chimera 14 and 15, Fig. 6d) abrogated interaction with HvSRF3^CD^ (Fig. 7e). Interestingly, of all 11 MLA13 contact residues in AVR_A13_, only five residues abrogate MLA13 activation upon exchange to alanine ^46^. All five of these ‘AVR_A13_ avirulence residues’ are located within the AVR_A13_ central loop region required for interaction of AVR_A13_ with HvSRF3 (‘AVR_A13_ virulence residues’). We therefore hypothesized that the AVR_A13_ avirulence residues overlap with AVR_A13_ virulence residues. To test this, we determined the ability of the here-generated chimeras to activate MLA13. For this, we co-expressed the constructs in *N. benthamiana* leaves for macroscopic evaluation of MLA13-mediated cell death. Additionally, we expressed the constructs in barley protoplasts to quantitatively assess MLA13-mediated cell death by determining the reduction of luciferase reporter activity as a proxy for cell viability^14^. As expected, CSEP0371 did not activate MLA13 cell death, and only the chimeras that can bind HvSRF3 activated MLA13 to the same extent as native AVR_A13_ (Fig. 7f and 7g, Supplementary Fig. 11c). Notably, with respect to chimera 12 and chimera 14, the exchange of the amino acids E44 and H45 of AVR_A13_ to R44 and D45 of CSEP0371 appear to impair the activation of MLA13 and abrogates the association with HvSRF3, underlining a central role of these residues in AVR_A13_ avirulence and presumed virulence function on HvSRF3 (Fig. 7f and 7g). In agreement with this hypothesis, neither of the AVR_A13_ variants (AVR_A13_^Y52A/G60A^, AVR_A13_^Y52A/F65A^, AVR_A13_^Y52A/G60A/F65A^)^46^ that were previously shown to have lost avirulence activity (activation of MLA13 cell death)^46^ was able to bind HvSRF3-L1^CD^ or HvSRF3-L2^CD^ (Fig. 7h, Supplementary Fig. 11d) or caused the previously observed (Fig. 4h) AVR_A13_-mediated changes to HvSRF3 PM abundance (Supplementary Fig. 12a to 12c).

**Figure 7:**
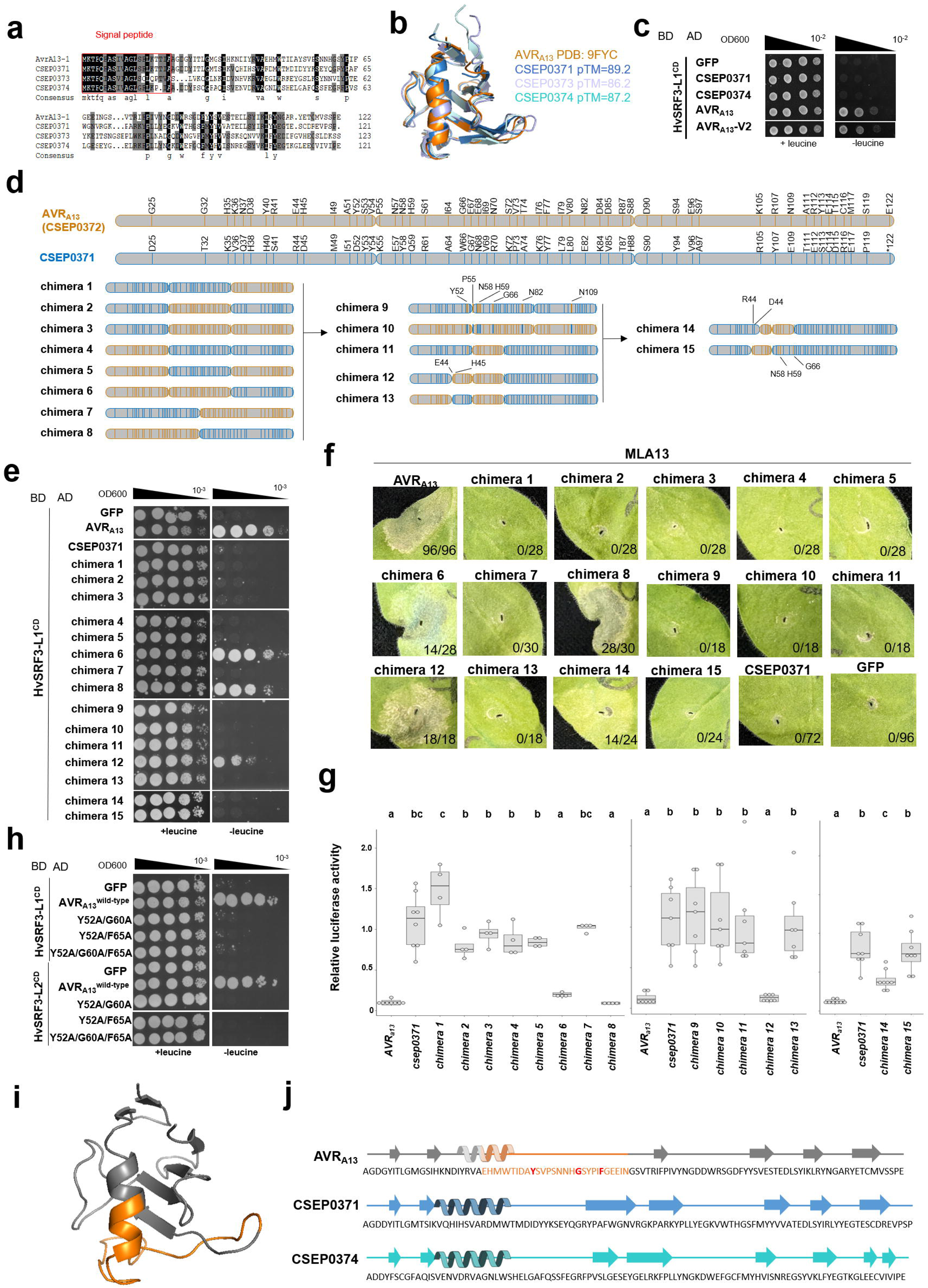
A central loop-containing region in AVR_A13_ specifies HvSRF3 interaction and MLA13 activation. **a** Sequence alignment of *Blumeria hordei* CSEP protein family 34 members AVR_A13_ (CSEP0372), CSEP0371, CSEP0373 and CSEP0374. **b** Structural overlay of AVR_A13_ (CSEP0372, PDB 9FYC) with AlphaFold models of CSEP0371 (pTM=89.2), CSEP0373 (pTM=86.2) and CSEP0374 (pTM=87.2). **c** Yeast two-hybrid assay with cells transformed with B42 activation domain (AD)- and HA tag-fused constructs encoding GFP or effectors together with LexA binding domain (BD)-fused constructs encoding the cytoplasmic domain of HvSRF3 (HvSRF3^CD^). Growth of transformants was determined on selective growth medium with leucine (+leucine), and interaction of proteins was determined by leucine reporter activity reflected by growth of yeast on selective medium lacking leucine (-leucine). Figures shown are representatives of at least three independent transformation experiments and pictures were taken three days after drop out. **d** Chimeras composed of AVR_A13_ aa residues (orange) and CSEP0371 aa residues (blue) as indicated. Vertical lines indicate differential amino acid residues between AVR_A13_ and CSEP0371. **e** Yeast cells were transformed with B42 AD- and HA tag-fused constructs encoding GFP, AVR_A13_, CSEP0371 or chimeras together with HvSRF3-L1^CD^ fused to the LexA BD. Protein interaction was determined as described in c. **f** *GFP* or C-terminally *mYFP*-fused *AVR_A13_*, *CSEP0371* or *chimeras* were co-overexpressed with *Mla13-myc* in leaves of four-week-old *Nicotiana benthamiana* plants using *Agrobacterium*-mediated transformation. Cell-death was determined three days post transformation. **g** Mesophyll protoplasts of barley cultivar Golden Promise were co-transfected with a *pZmUBQ:luciferase* reporter plasmid, *Mla13,* and constructs carrying *GFP*, *CSEP0371*, *AVR_a13_* or *chimeras* all in *pIPKb002*. Luciferase activity was determined at 16 h post transfection and normalized to the respective *GFP* sample. Values were obtained from at least four independent transfections performed on at least three independent days. Differences between samples processed simultaneously were assessed by analysis of variance (ANOVA p ≤ 2.3e-06) and subsequent Tukey post hoc test. Samples marked by identical letters in the plots do not differ significantly (p>0.05) in the Tukey test. **h** Yeast cells were transformed with B42 AD- and HA tag-fused constructs encoding GFP, AVR_A13_ wild type or its mutants together with HvSRF3-L1^CD^ or HvSRF3-L2^CD^ fused to the LexA BD. Protein interaction was determined as described in c. Signal peptides were depleted from all effector-encoding constructs (c, e, f, g, h). **i** AVR_A13_ structure (PDB 9FYC) indicating the central region (aa 44-70) required for HvSRF3 binding and MLA13 activation. **j** Structural arrangement of AVR_A13_ (PDB 9FYC), CSEP0371 (pTM=89.2) and CSEP0374 structure (pTM=87.2). Central region required for HvSRF3 binding and MLA13 activation in orange (i, j).

Together with the direct recognition of AVR_A13_ by MLA13^14,46^, our data suggests that HvSRF3 does not play a role for MLA13-mediated recognition of AVR_A13_. To test this genetically, we determined AVR_A13_-mediated activation of MLA13 in the barley *srf3* lines. In accordance with our hypothesis, MLA13-mediated cell death was quantitatively similar between wild-type and both homozygous *srf3* lines (Supplementary Fig. 12d). We conclude that the central region encoded by AVR_A13_ aa E44 to aa N70 (Fig. 6i and 6j) is required for direct activation of MLA13 and interaction with the host target HvSRF3 and therefore for an SRF3-mediated virulence function of the effector. It is thus plausible that MLA13 has evolved specifically to detect the region of AVR_A13_ required to inhibit HvSRF3 function.

## Discussion

We report the interactomes of five RNase-like *Bh* AVR_A_ effectors in the barley host. Despite their structural homology, individual AVR_A_ effectors engage with distinct sets of host proteins (Fig. 1, Supplementary Data 1). These findings experimentally support the hypothesised neo-functionalization of *Bh* RALPH effectors and align with previous studies that independently identified variable host targets for RALPH family members^10^. By deeper investigation of the most conserved *Bh* avirulence effector AVR_A13_, we demonstrate virulence activity and association with the barley RLK SRF3, for which the KD was shown to encode binding specificity (Fig. 2 and Fig. 3). Enhanced *Bh* proliferation on barley *srf3* mutants and *AVR_a13_* barley lines suggests that this interaction is indeed of biological relevance but does not exclude that in addition to HvSRF3, AVR_A13_ also targets other host proteins for promoting fungal infection^47^. Notably, the HvSRF3-binding activity of AVR_A13_ cannot be performed by AVR_A_s that are recognized by other *Mla* alleles (Fig. 1 and Fig. 3a), suggesting variable virulence functions for individual AVR_A_ effectors. Collectively, our results suggest that pathogen effectors that have originated from the same species and have substantially contributed to the diversification of host NLR receptors, do not necessarily converge on the same host targets for virulence.

Similar to the *Bh* AVR_A_ RALPH effectors, the *Magnaporthe oryzae* AvrPik effectors recognized by allelic variants of Pik1 are structural homologs^48–50^. Pik1 detects allelic AvrPik variants through the integrated Heavy Metal–Associated (HMA) domain, and the allelic AvrPik effectors exert virulence activity by binding the rice heavy metal–associated isoprenylated proteins (HIPPs) OsHIPP19 and OsHIPP20^50,51^. This interaction is conserved across all tested AvrPik variants, suggesting that Pik1 evolved to recognize AvrPik virulence activity by mimicking HIPP19 and HIPP20 HMA domains ^51^. In turn, the divergent *M. oryzae* effectors AVR-Pia, AVR1-CO39 and Pwl2 cannot bind HIPP19 and HIPP20, and PWL2 instead associates with OsHIPP43^51,52^. This prompts us to speculate that *Bh* AVR_A_ and potentially other RALPH effectors may likewise converge on specific domains within distinct barley proteins to promote *Bh* virulence. We show that the specificity of AVR_A13_ association with HvSRF3 is determined by the KD of HvSRF3, suggesting that KD-containing proteins that participate in diverse physiological processes may serve as convergent targets for AVR_A_ effectors. Even within the AVR_A13_ effector family, HvSRF3 associates only with AVR_A13_, despite CSEP0371 and CSEP0374 being the most closely related *Bh* effector proteins in terms of sequence and structure (Fig. 7)^8^. This observation is significant because prior structural analyses of AVR_A_ effectors led to the hypothesis that members of the same RALPH subclass share conserved virulence activities^18^. It will be interesting to determine whether CSEP0371 and CSEP0374 interact with other HvSRFs or target distinct KD-containing proteins.

The association specificity of AVR_A13_ with HvSRF3 is determined by the HvSRF3 KDs, which appear to be pseudokinases (Fig. 3). Similarly, no kinase activity has been detected for AtSRF3 *in vitro*^39^, and the SRF member AtSUB/AtSRF9 appears to carry a catalytically inactive kinase domain^28,30,31^. Despite this, SRF3 and SUB KDs are required for SRF-specific phenotypes^28,30,31,39^. Based on the interaction of HvSRF3 and HvBAK1, we speculate that HvSRF3 might rely on HvBAK1 for signaling. This is consistent with multiple independent studies involving different SRF family members^35–37^. However, we cannot exclude that similar to the kinase-inactive LRR-RLKs BAK1-INTERACTING RECEPTOR-LIKE KINASE 1 and 2 (BIR1 and BIR2)^53,54^, HvSRF3 acts as a general regulator of HvBAK1 activity and that AVR_A13_ positively regulates these processes. Yet, this scenario appears unlikely because BAK1 is a well-established positive regulator of plant immunity, and HvSRF3-L1/HvSRF3-L2 do not influence flg22-induced ROS production (Supplemental Fig. 10), a response that fully depends on BAK1^55,56^. Either way, the AVR_A13_-mediated inhibition of the HvSRF3-HvBAK1 association suggests that this interaction between HvSRF3 and HvBAK1 is indeed of biological relevance. This is also notable with respect to numerous reports demonstrating that tandem kinase genes mediate resistance to biotrophic fungi in barley and wheat^57^. Tandem kinase effector recognition seems to induce conformational changes within paired KDs^57–59^. This is relevant, as tandem kinases typically pair an active kinase with a pseudokinase^59–65^, a configuration conceptually similar to our observations, where AVR_A13_ binding to the HvSRF3 pseudokinase domain influences its association with the active HvBAK1 kinase. Tandem-kinase-recognized effectors may thus enhance fungal virulence by targeting immune RLK complexes with active and inactive kinases (e.g., BAK1/SERKs and SRFs).

*AVR_a13_*-expressing *A. thaliana* lines mimic *srf3* mutants under low iron conditions (Fig. 5), suggesting the conservation of the AVR_A13_ function on SRF3 between dicots and monocots and underlining negative regulation of SRF3 by AVR_A13_^39^. *A. thaliana srf3* mutants exhibit increased iron levels under sufficient iron conditions. In turn, AtSRF3 is involved in the upregulation of low iron response genes under iron deficient conditions^39^. As such, SRF3 responds to differential iron levels and regulates iron homeostasis. Importantly, SRF3-mediated upregulation of low iron response genes is impaired during PTI responses^39^.

The regulation of iron homeostasis upon microbial attack is crucial for all hosts. Reducing the iron content in the sites of infection to deprive the pathogens of iron (nutritional immunity) and also the over-accumulation of redox-active iron at the infection sites are effective host defense strategies^66^. Both, intracellular ferrous iron depletion and accumulation of extracellular ferric iron early during powdery mildew attack has been reported^41^, but it had remained unclear how iron influences fungal proliferation on the host. Here we found that reduced host iron availability has little effect on *Bh* proliferation, suggesting that withholding iron is not an effective *Bh* defense strategy. In turn, high levels of iron negatively regulate *Bh* disease development (Fig. 6, Supplementary Fig. 10a). *AVR_a13_*-expressing lines and *srf3* lines and are insensitive to this (Fig. 6). We thereby link both, SRF3 and AVR_A13_ function to iron-induced inhibition of fungal infection. Based on this, we speculate that SRF3 directly or indirectly conveys information about changes to iron levels in *Bh* infected cells and that this process i) might involve HvBAK1 and ii) is inhibited by AVR_A13_ to obstruct the resupply of immune active iron to the site of infection. Notably, it is entirely possible that AVR_A13_ targets SRF3 to indirectly manipulate other processes that are influenced by differential iron levels. Such processes may involve cell wall composition and integrity or oxidative stress responses^25,67,68^. Yet, we demonstrate that a putative SRF3-mediated regulation of the host redox status does not influence chitin-mediated ROS production.

AVR_A13_ binds HvSRF3 *via* a loop that also triggers MLA13-mediated cell death. Chimera and cryo-EM data show residues outside this loop are dispensable for MLA13 activation (Fig. 7)^46^, suggesting MLA13 specifically evolved to recognize the SRF3-interaction loop of AVR_A13_, while the MLA13 winged-helix domain enables general RNase-like effector binding. AVR_A13_’s overlapping avirulence–virulence function likely prevents *Mla13*-evading variants, accounting for its conservation in *Bh* isolates across decades^15,14^. In turn, virulent AVR_A13_-V2 does not activate MLA13-mediated cell death but carries the shared HvSRF3-binding and MLA13-activity loop region. We have previously demonstrated that AVR_A13_-V2 maintained the residues for MLA13 interaction but blocked MLA13-mediated cell death and have thereby described a rare pathogen-derived mechanism for escaping NLR-mediated resistance^23^. The data presented here can explain that this complex mechanism allows the escape from NLR-mediated resistance whilst maintaining overlapping virulence propensities, which here appears superior to the escape from NLR-binding^14^.

Enhanced fungal virulence on barley *srf3* lines together with the overlap between HvSRF3-binding residues of AVR_A13_ and those required for MLA13 activation, highlight SRF3’s central role in plant immunity. This is also supported by independent lines of evidence: (i) the involvement of SRF3 in autoimmunity mediated by the DM2h NLR in *A. thaliana* ecotype L*er*^69,70^, suggesting that AtSRF3 acts as a decoy for an unknown effector^71^; (ii) the regulation of nutritional immunity by AtSRF3 in *A. thaliana* roots^39^; and (iii) the targeting of rice SRF3 by an effector from a smut fungus^37^. Taken together, these findings highlight SRF3 as a conserved and pivotal regulator of plant-microbe interactions, with roles that extend well beyond *Bh* infection in barley. The ability of AVR_A13_ to also bind AtSRF3 will provide us with a genetically tractable system to further dissect SRF3 function during pathogen attack.

## Methods

### Plant material and growth

Barley was grown at 19°C, 70% relative humidity, and under a 16 h photoperiod. *Nicotiana benthamiana* was grown as described^72^. *Arabidopsis thaliana* plants were grown at 21 °C, under an eight-hour photoperiod for experimental approaches or a 16-hour photoperiod for seed propagation.

### *Blumeria hordei* maintainance and infections

Maintenance of *Blumeria hordei* isolate K1 was carried out as described previously^15^. For fungal propagation and infection assays, one-week-old barley plants were transferred to an incubator with *Bh* isolate K1 infected barley cultivar Golden Promise incubator and thereby inoculated. For iron treatment, plants were sprayed with water containing 0.1% (v/v) Tween 20 (mock) or with a 0.5 g/L FeEDDHA solution containing 0.1% (v/v) Tween 20 and thereafter transferred to the *Bh* incubator. To quantify infection, the adaxial and abaxial surfaces of the first leaf were photographed, and *Bh* colonies were counted five days post-inoculation.

### *Blumeria hordei* infection and quantification on *in vitro* grown barley

For *in vitro* growth, barley seeds were surfaces-sterilized in 6% sodium hypochlorite, shaking for one hour, washed five times with sterile Milli-Q water and germinated on wet sterile Whatman filter paper. Four-day-old germinated seedlings were transferred into Weck glasses containing sterile ½ MS medium with either 10 µM, 100 µM or 250 µM FeNaEDTA. Except for FeNaEDTA, the MS medium was prepared according to the Duchefa recipe (M0255). Plants were grown for another three days prior to infection with *Blumeria hordei* (*Bh*) isolate K1. At seven days post-infection, leaf tissue from three independent plants was pooled and frozen in liquid nitrogen. Tissue was ground to a fine powder using a Retsch Beat Beater (2 × 1 min at 30 Hz), and total DNA was extracted according to Edwards & Thompson (1991)^73^. Fungal biomass was quantified by qPCR using the GoTaq® qPCR kit containing 2X BRYT Green® dye master mixes on the Bio-Rad CFX Connect Real-Time PCR System. *Bh* GAPDH primers (5’GGAGCCGAGTACATAGTAGAGT’3, 5’GGAGGGTGCCGAAATGATAAC’3; GenBank: CAUH01004767) and barley GAPDH primers (5’TTTGGTATTATTGAGGGTCTGATGA’3, 5’TGCTGCTGGGGATGATGTT’3; GenBank: X60343.1) were used. Data were visualized and processed in Bio-Rad CFX Manager. Fold changes were calculated using the delta-delta-Ct method after normalization of the *Bh* GAPDH Ct values to the Ct mean of the barley GAPDH housekeeping gene.

### Generation of expression constructs

*SRF3* variants, *CSEP00371, CSEP00374*, and the *AVR_a13_-CSEP00371 chimeras* were either amplified from cDNAs of *A. thaliana* Col-0 or barley cv. Golden Promise using Q5 High-Fidelity DNA Polymerase (NEB) or synthesized from GeneArt (Thermo Scientific) as pDONR221 entry clones or with overhangs for restriction/ligation into Golden Gate Level 0 vectors^74^. For stable transformation of barley, synthesized *BirA-4myc* and *AVR_a_-BirA-4myc*^23^ constructs were transferred from donor vectors into the expression vector pIPKb002^75^. The generation of *pAtUBI10:FLAG-AVR_a_* constructs for expression of *AVR_a13_* in *A. thaliana* was described previously^15^. For determination of HvSRF3 localization in barley, HvSRF3 constructs were cloned together with *pZmUbi* promoter (derived from the pIPKb002 vector)^75^ and a C-terminal mYFP tag derived from the pXCSG-GW-mYFP vector^76^ into pICH47732^74^ by restriction enzyme digest/ligation. For Y2H interaction studies, respective genes were either described previously ^14^ or were transferred from gateway entry or donor vectors into the expression vectors pLexA-GW, or pB42AD-GW^14^ using LR Clonase II (Thermo Scientific). For split-luciferase assays, respective genes were cloned into expression vector pCAMBIR1300-nLUC or pCAMBIR1300-cLUC by NEBuilder HiFi DNA Assembly as described^24^. For CoIP assays, SRF3 constructs, the cauliflower mosaic virus 35S promoter sequence and a 6HA or GFP tag sequences were cloned into expression vector pICH47732^74^ using restriction enzyme digestion/ligation or NEBuilder HiFi DNA Assembly. AVR_A13_-mYFP constructs were described previously^14^. For BiFC assays, respective constructs were cloned into expression vector pICH47732 with C-terminal YFP^N^ (NE) or YFP^C^ (CE) by NEBuilder HiFi DNA Assembly^74,77^. For subcellular localization studies, respective genes were cloned together with the cauliflower mosaic virus 35S promoter and an mCherry or GFP tag into expression vector pICH47732^74^ using restriction enzyme digestion/ligation or NEBuilder HiFi DNA Assembly. For cell death assays in *N. benthamiana* leaves GFP, AVR_A13_, CSEP0371 or chimera-encoding constructs were cloned together with the cauliflower mosaic virus 35S promoter and a C-terminal mYFP-tag derived from the pXCSG-GW-mYFP vector^76^ into pICH47732^74^ by restriction enzyme digest/ligation. The *35S:Mla13* construct was generated previously^14^. For cell death assays in barley protoplasts, GFP, CSEP0371 or chimera-encoding constructs were cloned from pDONR221 into pIPKb002^75^ using LR Clonase II (Thermo Scientific). *pZmUbi:luciferase*, *Mla13* (in pIPKb002) and *AVR_a13_* (in pIPKb002) constructs were generated previously^14,23^.

### Generation of cas9 and gRNA constructs

Twenty-nucleotide target sequences present immediately adjacent to a Protospacer Adjacent Motif (PAM) were selected using the CRISPOR web server (https://crispor.gi.ucsc.edu/) and verified using the RNAfold web server (http://rna.tbi.univie.ac.at/cgi-bin/RNAWebSuite/RNAfold.cgi) as described^78^ for the *HvSRF3* genes. Two single guide RNAs (gRNAs) were selected (g1 5’-GCAAACGGCGGAGACCCCTG-3’ and g2 5’-AATCGCCAAAGCTTCCCGGG-3’) and generated using the CasCADE modular vector system, which employs a hierarchical Golden Gate cloning strategy^79^. Respective oligo target sequences and BsaI restriction sites were annealed and cloned into BsaI–linearized vectors pIK1 to pIK2 containing the *OsU3* promoter or pIK5 to pIK6 containing the *TaU6* promoter. The resultant vectors were digested with the Esp3I enzyme and assembled into the pIK60 vector. To combine the two gRNA fragments with the *cas9* expression unit (pIK83) and an auxiliary unit vector (pIK155) into vector pIK48, a BsaI restriction digest followed by ligation was used. Last, all expression units were mobilized into the binary vector p6-d35S-TE9 (DNA Cloning Service) *via* SfiI restriction digest.

### Generation of transgenic plants

The floral dip method^80^ was used to generate transgenic *A. thaliana* plants and homozygous single copy transgenic lines were generated base on segregation. Transgenic barley lines overexpressing *BirA* and *BirA* fusions were generated by *Agrobacterium tumefaciens*– mediated transformation of immature embryos derived from *Hordeum vulgare* ssp. *vulgare* (barley) cultivar Golden Promise Fast^81^ as described previously^82^. For this, the strain *A. tumefaciens* AGL1 carrying pIPKb002 vectors with *BirA-4myc* or respective *AVR_A_-BirA-4myc* constructs were used^23^. Transformants were selected on hygromycin and the presence of transgenes was confirmed by PCR using a pIPKb002 forward (5′-CACCCTGTTGTTTGGTGTTACTTC -3′) and *BirA*-specific reverse primers (5′-TCGATCACAGGAAGAACAGCAAC-3′) according to Edwards & Thompson (1991)^73^. Effector protein expression was determined by anti-Myc Western blot analysis of respective leaf extracts. T2 (*AVR_a13_-BirA-4myc* #31-1 and #44-6, BirA-4myc #32-1 and #32-2, *AVR_a9_-BirA-4myc* # 43-1 and #43-16) and respective T3 lines were generated (AVR_A13_-BirA-4myc #52 and #55, BirA-4myc #53 and #54) from independent transformants (T1). Barley CRISPR-Cas9 *srf3* lines were generated by *Agrobacterium tumefaciens*–mediated transformation of immature embryos derived from *Hordeum vulgare* ssp. *vulgare* (barley) cv. Golden Promise as described previously ^83^. For this, the strain *A. tumefaciens* AGL1 carrying the p6-d35S-TE9 vector with guide RNA1 (g2) and guide RNA2 (g2) were used (Supplementary Fig. 10d). Mutants were selected by PCR according to Edwards & Thompson 1991^73^ using target-side specific primers for the four *HvSRF3* genes (L1 fw: 5′-ATGTGGGTTGTGGCTTTTGC-3′ rv: 5′-GAGCCTGACATACATTGAGG-3′, L2 fw: 5′-CGTTCCTAATGGCTTATTTG-3′ rv: 5′-GCCGTGAAGTTTCCTAGGCT-3′, L3 fw: 5′-GCACTCGCTGACCTACACTT-3′ rv: 5′-TGTTTGGAAGGTGCCCACTT-3′, L4 fw: 5′-CTCCTATGGCTCGCCTTCTG-3′ rv: 5′-TCCCGGTTAGCCTGTTGTTC-3′) followed by sanger sequencing of the amplicons and the absence of the Cas9 transgene was confirmed by PCR (fw: 5′-CCTGCCTTCATACGCTATTTATTTG-3′ rv: 5’-TTAATCATGTGGGCCAGAGC-3’)

### Determination of late root growth responses

For surface sterilization, Col-0 and *AVR_a13_*-expressing *A. thaliana* seeds that had been produced under uniform growth conditions at the same time were placed for one hour in opened 1.5-mL tubes in a sealed box containing chlorine gas generated from 130 mL of 10% sodium hypochlorite and 3.5 mL of 37% hydrochloric acid. Seeds were then sown on plates containing KH_2_PO_4_ at 100 µM, MgSO_4_ 7H_2_O at 100 µM, K_2_SO at 25 µM, CaCl_2_ 2xH_2_O at 25 µM, KCl at 50 µM, H_3_BO_3_ at 30 µM, MnSO_4_, H_2_O at 5 µM, ZnSO_4_,5H_2_O at 1 µM, CuSO_4_,5H_2_O at 1 µM, (NH_4_)_6_Mo_7_O_24_ at 0.1 µM, NH_4_NO_3_ at 1 mM, Na-Fe-EDTA at 50 µM, MES at 250 µM and Agar type A (A4550, Sigma Aldrich) at 0.8% with pH adjusted at 5.7 with KOH. Stratification was performed in the dark at 4 °C for 2-3 days. Five days after growth under standard condition, plates were transferred to identical media described above, either with Iron (50 µM FeEDTA, +Fe) or without iron (0 µM Na-Fe-EDTA, + 100 µM Ferrozine, -Fe). For this, six plants per genotype were transferred to four 12 × 12-cm plates in a block design with genotypes alternating across the plates (top left, top right, bottom left, and bottom right). After transfer, the plates were scanned every day for three days using the EPSON flatbed scanners (Perfection V600 Photo, Seiko Epson CO., Nagano, Japan). Quantification was performed by stacking the images of each day using a macro in Fiji (Macro_Match_Align) and the root length for each day per genotype in each condition to evaluate the root growth rate in Fiji using the segmented line. Then, the mean root growth rate for each day and line was calculated. These values were used to calculate the root growth rate per day for each plant. Then, the root growth rate was divided per day under -Fe for each plant by the mean root growth rate per day to the +Fe media for the entire related genotype. This ratio was used as the late root growth response to low iron levels as described^39^.

### PAMP-induced reactive oxygen species production assay

Leaf discs (0.38 cm^2^) from 10-day-old barley seedlings were floated on 200 µl water in a white 96-well plate (one disk/well) overnight. The water was replaced with 100 μl of a solution containing 10 μM L-012 (Wako chemicals) and 1 μg/ml of horseradish peroxidase (SIGMA) supplemented with 100 nM flg22 or 10 µg/ml chitin to elicit the production of ROS. Luminescence was measured on a, Infinite M200 PRO plate reader (Tecan) for one second/well/minute over a time course of 30 minutes.

### Confocal microscopy

Fluorescent signals were visualized by laser scanning confocal microscopy (LSCM) using a TCS SP8 confocal microscope (Leica). GFP signal was excited with an Argon laser at a wavelength of 488 nm and emitted light was detected at 500–520 nm. mYFP signal was elicited with an Argon laser at a wavelength of 514 nm and detected at 520–540 nm. mCherry signal was excited with a 561 DPSS laser at a wavelength of 561 and emission was detected at 590 to 630 nm.

### Proximity-dependent protein labelling using the BirA/biotin system

Barley plants were grown a 16-hour light/ eight-hour dark photoperiod for four weeks. Leaves were then infiltrated with 250 µM biotin solution in 10 mM MgCl₂. After 36 h, leaf samples (300 mg per tube with 2 × 5 mm glass beads) were collected and immediately frozen in liquid nitrogen. For Strep-Tactin enrichment, frozen leaf samples were ground three times for one minute at 20 Hz with samples returned to liquid nitrogen between grinding cycles. SDT lysis buffer (prepared as 10 mL 10% SDS, 2.5 mL 1 M DTT, 2.5 mL 1 M Tris-HCl, pH 7.5) was pre-heated to 95°C. A total of 550 µL of hot SDT buffer was added to each frozen powder sample, immediately inverted and vortexed. Samples were incubated for 5 min at 95°C and briefly centrifuged at 2000 × g, followed by sonication for 10 min. Lysates were centrifuged for 10 min at 13000 × g, and 500 µL of the supernatant was transferred to a fresh tube. A 10 µL aliquot was diluted with 40 µL PBS and mixed with 12.5 µL 5× SDS loading buffer for western blot analysis (total protein). To deplete biotin, 666 µL methanol was added to 500 µL supernatant, followed by 166 µL chloroform. Samples were vortexed and then mixed with 300 µL Milli-Q water. After centrifugation for 10 min at 2000 × g, both upper and lower phases were discarded, and the pellet was retained. The pellet was washed twice with 600 µL methanol, each time vortexed and sonicated for 10 min to break up the pellet, then centrifuged for 10 min at 13000 × g. Supernatant was completely removed and the pellet was air-dried for up to 5 min. The pellet was resuspended in 500 µL SDT buffer, vortexed, sonicated for 10 min, and vortexed again. Samples were incubated for 30 min at room temperature with shaking on an Eppendorf Thermomixer at 1000 rpm, then diluted to 4 mL in PBS buffer to achieve a final SDS concentration of 0.5%. A 50 µL aliquot was mixed with 12.5 µL 5× SDS loading buffer for western blot analysis (biotin-depleted sample). Strep-Tactin beads (50 µL slurry per sample Thermo Fisher, 20347) were washed three times with 1 mL PBS (centrifugation for 2 min at 2000 × g) and resuspended in 100 µL PBS. Fifty microliters of bead slurry were added to each protein sample, and the mixture was incubated overnight at 4°C in a 15 mL tube with gentle mixing. Beads were collected by centrifugation for 3 min at 2000 × g, and 50 µL of the supernatant was taken for western blot analysis (supernatant fraction). The beads were then washed with 2 mL PBS (pH 7.2) containing 2% SDS, centrifuged for 3 min at 2000 × g, and the supernatant was removed. The wash step was repeated six times with 10 mL PBS. Washed beads were resuspended in 1 mL PBS, transferred to a 1.5 mL microcentrifuge tube, and centrifuged for 3 min at 2000 × g. Beads were finally resuspended in 200 µL PBS. 20 µL of bead slurry was mixed with 20 µL of 2× SDS loading buffer containing 20 mM biotin. Samples were centrifuged for 3 min at 2000 × g, and the supernatant was carefully removed. The remaining beads were stored at −80°C and subsequently subjected to Liquid Chromatography-Mass Spectrometry analysis.

### Liquid Chromatography-Mass Spectrometry

Proteins from Strep-Tactin beads were submitted to an on-bead digestion. Beads were resuspended in 25 µL digestion buffer 1 (50 mM Tris-HCl, pH 7.5; 2 M urea; 1 mM DTT; 5 ng/µL trypsin) and incubated for 30 min at 30°C in a Thermomixer (400 rpm). Beads were then pelleted by centrifugation, and the supernatant was transferred to a fresh tube. Next, 50 µL digestion buffer 2 (50 mM Tris-HCl, pH 7.5; 2 M urea; 5 mM CAA) was added to the beads. After mixing, the beads were pelleted again, and the supernatant was combined with the previous fraction. The combined supernatants were incubated overnight at 32°C in a Thermomixer (400 rpm) protected from light. Digestion was terminated by adding 1 µL trifluoroacetic acid (TFA), and peptides were desalted using C18 Empore disk membranes following the StageTip protocol ^85^. Dried peptides were re-dissolved in 2% ACN, 0.1% TFA (10 µL) and diluted to 0.1 µg/µL for analysis. Samples were analyzed using an EASY-nLC 1200 (Thermo Fisher Scientific) coupled to a Q Exactive Plus mass spectrometer (Thermo Fisher Scientific). Peptides were separated on 16 cm frit-less silica emitters (New Objective, 75 µm inner diameter), packed in-house with reversed-phase ReproSil-Pur C18 AQ 1.9 µm resin. Peptides were loaded on the column and eluted for 115 min using a segmented linear gradient of 5% to 95% solvent B (0 min : 5%B; 0-5 min -> 5%B; 5-65 min -> 20%B; 65-90 min ->35%B; 90-100 min -> 55%; 100-105 min ->95%, 105-115 min ->95%) (solvent A 0% ACN, 0.1% FA; solvent B 80% ACN, 0.1%FA) at a flow rate of 300 nL/min. Mass spectra were acquired in data-dependent acquisition mode with a TOP15 method. MS spectra were acquired in the Orbitrap analyzer with a mass range of 300–1750 m/z at a resolution of 70000 FWHM and a target value of 3×10^6^ ions. Precursors were selected with an isolation window of 1.3 m/z. HCD fragmentation was performed at a normalized collision energy of 25. MS/MS spectra were acquired with a target value of 10^5^ ions at a resolution of 17500 FWHM, a maximum injection time (max.) of 55 ms and a fixed first mass of m/z 100. Peptides with a charge of +1, greater than 6, or with unassigned charge state were excluded from fragmentation for MS^2^, dynamic exclusion for 30s prevented repeated selection of precursors. Alternatively, Samples were analyzed using an Ultimate 3000 RSLC nano (Thermo Fisher) coupled to an Orbitrap Exploris 480 mass spectrometer equipped with a FAIMS Pro interface for Field asymmetric ion mobility separation (Thermo Fisher). Peptides were pre-concentrated on an Acclaim PepMap 100 pre-column (75 µM x 2 cm, C18, 3 µM, 100 Å, Thermo Fisher) using the loading pump and buffer A** (water, 0.1% TFA) with a flow of 7 µL/min for 5 min. Peptides were separated on 16 cm frit-less silica emitters (New Objective, 75 µm inner diameter), packed in-house with reversed-phase ReproSil-Pur C18 AQ 1.9 µm resin (Dr. Maisch). Peptides were loaded on the column and eluted for 130 min using a segmented linear gradient of 5% to 95% solvent B (0 min : 5%B; 0-5 min -> 5%B; 5-65 min -> 20%B; 65-90 min ->35%B; 90-100 min -> 55%; 100-105 min ->95%, 105-115 min ->95%, 115-115.1 min -> 5%, 115.1-130 min ->5%) (solvent A 0% ACN, 0.1% FA; solvent B 80% ACN, 0.1%FA) at a flow rate of 300 nL/min. Mass spectra were acquired in data-dependent acquisition mode with a TOP_S method using a cycle time of 2 seconds. For field asymmetric ion mobility separation (FAIMS) two compensation voltages (- 45 and -60) were applied, the cycle times for both compensation voltages were set to 1 second. MS spectra were acquired in the Orbitrap analyzer with a mass range of 320–1200 m/z at a resolution of 60 000 FWHM and a normalized AGC target of 300%. Precursors were filtered using the MIPS option (MIPS mode = peptide), the intensity threshold was set to 5000, Precursors were selected with an isolation window of 1.6 m/z. HCD fragmentation was performed at a normalized collision energy of 30%. MS/MS spectra were acquired with a target value of 75% ions at a resolution of 15,000 FWHM, at an injection time of 120 ms and a fixed first mass of m/z 120. Peptides with a charge of +1, greater than 6, or with unassigned charge state were excluded from fragmentation for MS^2^. The mass spectrometry proteomics data have been deposited with the ProteomeXchange Consortium via the PRIDE partner repository under the dataset identifier PXD07175^86^. Raw data were processed using MaxQuant software (version 1.6.3.4, http://www.maxquant.org/) with label-free quantification (LFQ) and iBAQ enabled^87,88^. MS/MS spectra were searched by the Andromeda search engine against a combined database containing the sequences from barley (Barley_Morex_V2_gene_annotation_PGSB.all.aa^89^) and sequences of 248 common contaminant proteins and decoy sequences. Trypsin specificity was required and a maximum of two missed cleavages allowed. Minimal peptide length was set to seven amino acids. Carbamidomethylation of cysteine residues was set as fixed, oxidation of methionine and protein N-terminal acetylation as variable modifications. Peptide-spectrum-matches and proteins were retained if they were below a false discovery rate of 1%. Statistical analysis of the MaxLFQ values was carried out using Perseus (version 1.5.8.5, http://www.maxquant.org/). Quantified proteins were filtered for reverse hits and hits “identified by site” and MaxLFQ values were log2 transformed. After grouping samples by condition only those proteins were retained for the subsequent analysis that had 2 valid values in one of the conditions. The relative amounts of each identified protein across samples were quantified by using label-free quantification (LFQ), and protein abundances within a sample were quantified using intensity-based absolute quantification (iBAQ)^88^. The data were analyzed by a t-test-based analysis (LFQ data) and volcano plot analysis (iBAQ data). For t-test-based analysis the enrichment of proteins between wild-type or BirA control and the AVR_A_-BirA samples, we used the raw data (≥2 valid LFQ values) to calculate two-sample t-tests with a permutation-based false discovery rate (FDR) of 5%. The enriched proteins were designated as interactors of the AVR_A_s, respectively, if they had a p-value <0.05 and log2 fold change >1 over the wild-type and BirA-4myc controls.

For volcano plot analysis, quantified proteins were grouped by condition and only those hits were retained that had 3 valid values in one of the conditions. Missing values were imputed from a normal distribution (1.8 downshift, separately for each column). Volcano plots were generated in Perseus using an FDR of 5% and an S0=1. The Perseus output was exported and further processed using excel (Supplementary Data 1).

### Barley protoplast-based cell death assay

Quantification of cell death of barley mesophyll protoplasts using luciferase activity as a proxy for cell viability was performed as described^14^ with the following three modifications: 1. A GFP-encoding construct in pIPKb002 was used instead of the empty vector. 2. For each transfection of 300 µL of protoplasts, the plasmid solutions with a total volume of 30 µL containing 4 to 5 µg of the *luciferase* reporter plasmid, 10 µg of the *Mla13* construct, and 5 to 10 µg of the GFP or the respective effector construct. 3. Luciferase activity was determined at 1 seconds/well using a Tecan Plate Reader (TECAN, Infinite® 200 PRO).

### Agrobacterium-mediated transformation of Nicotiana benthamiana leaves

*Agrobacterium tumefaciens* GV3101 pMP90K was freshly transformed by electroporation with the respective constructs of interest, unless otherwise specified. For each construct, 10 ml of Luria-Bertani (LB) Lennox containing appropriate antibiotics were inoculated with three single colonies for each construct, and cultured over night at 28°C, 200 rpm shaking to a maximum OD_600_ = 1.5. Bacteria were harvested by centrifugation at 2500 × *g* for 10 min and resuspended in infiltration buffer (10 mM MES, pH 5.6, 10 mM MgCl_2_, and 200 mM acetosyringone) to the desired final OD_600_ (OD_600_ = 1 unless otherwise stated). Bacterial suspensions were incubated two to four hours at 28°C at 180 rpm, mixed equally and subsequently infiltrated into the adaxial site of leaves of three to four-week-old *N. benthamiana* plants with a needleless syringe.

### Cell death assay in *Nicotiana benthamiana* leaves

*A. tumefaciens* carrying *Mla13* or the silencing inhibitors *p19* or *CMV2b* were not freshly transformed but streaked from glycerol stocks to LB Lennox agar plates and grown as described above. Suspensions of *Mla13, p19* and *CMV2b*-carrying bacteria was adjusted to OD_600_ = 2, while bacteria freshly transformed with effector constructs were adjusted to an OD_600_ = 2.4. Bacterial suspensions were then mixed at equal ratios and infiltrated into *N. benthamiana* leaves. At three days post-infiltration cell death phenotypes were assessed, and leaves were harvested for protein expression analysis.

### Co-Immunoprecipitation assay

*Nicotiana benthamiana* leaf tissue (300 mg) co-expressing the constructs of interest was harvested three days post infiltration and placed in two mL safe-lock tubes containing two 5 mm glass beads and immediately frozen in liquid nitrogen. Frozen tissue was ground to a fine powder using a Retsch Beat Beater (2 × 1 min at 30 Hz) and thereafter resuspended in 1.5 mL ice-cold protein lysis buffer (150 mM Tris-HCl, pH 7.5; 150 mM NaCl; 10% glycerol; 2 mM EDTA; 6 mM DTT; 0.5% Triton X-100; 1 mM PMSF) supplemented with cOmplete™ Protease Inhibitor Cocktail (Roche). Samples were rotated at 4°C for 30 min to extract total proteins. Protein extracts were centrifuged twice at 16,000 × g for 10 min, and the supernatant was collected as the input fraction. An aliquot of the input was retained for immunoblot analysis. For immunoprecipitation, 5 µL of washed GFP-Trap magnetic beads (Chromotek) were added to each protein extract and incubated on a rolling wheel at 4°C for 3h. Beads were magnetically separated and washed three times with 1 mL wash buffer (150 mM Tris-HCl, pH 7.5; 150 mM NaCl; 10% glycerol; 2 mM EDTA). Bound proteins were eluted by boiling the beads in 2x SDS loading buffer for subsequent immunoblotting.

### Split-luciferase complementation (split-LUC) assay

Split-LUC assays in *N. benthamiana* were performed as described^24^. For this, *N. benthamiana* leaves expressing the constructs of interest were harvested three days post-infiltration and sprayed with 1 mM D-luciferin (Promega, E1602). After 10 min of incubation in the dark, the luminescence signals from leaves of at least three independent plants were detected using a CCD imaging system (ChemiDoc, Bio-Rad).

### Bimolecular Fluorescence Complementation (BiFC) assay

BiFC assays in *N. benthamiana* were performed as described^77^. *N. benthamiana* leaves expressing the constructs of interest were harvested two days post-infiltration and YFP complementation was determined by confocal laser scanning microscopy.

### Western blot analysis on plant-derived material

For analysis of protein levels and protein stability upon transgene expression, leaf material from barley, *A. thaliana* and *N. benthamiana* was harvested and frozen in liquid nitrogen. Leaf material was ground to a fine powder using pre-cooled adapters in a bead beater (Retsch) at 2 min, 30 Hz. At a ∼ 100 mg fresh weight / 300 µL buffer ratio^90^, ground tissue was submerged in plant protein extraction buffer (150 mM Tris-HCl, pH 7.5, 150 mM NaCl, 10 mM EDTA, 6 mM DTT, 10% (v/v) glycerol, 1 mM PMSF, 1.5% (v/v) plant protease inhibitor cocktail (Sigma)) and subsequently centrifuged at 16,000 × g at 4°C, 10 min. The supernatant was transferred to a fresh 2 ml tube and centrifuged at 4°C for 30 min, 16,000 × g. Extracts were diluted with 4x Laemmli buffer (Bio-Rad, 1610747) containing 10% β-mercaptoethanol and heated to 95°C for 10 min. Samples were centrifuged at 16,000 × g for 10 min prior to separation by SDS-polyacrylamide-gel electrophoresis (PAGE) (Bio-Rad), and blotted onto polyvinylidene fluoride (PVDF) membranes (Merck). Membranes were blocked in TBST with 3% non-fat milk and probed with either Strep-HRP (Sigma-Aldrich, S2438) or FLAG-HRP (Sigma-Aldrich, A8592), anti-GFP (abcam, ab6556), anti-LUC (Sigma-Aldrich, L0159), anti-HA (Sigma-Aldrich, 11867423001), anti-Myc (abcam, ab9106) or anti-mCherry (Thermo Fisher, 600-401-P16S) followed by anti-rabbit IgG-HRP (Santa Cruz Biotechnology, sc-2357), anti-rat IgG-HRP (Sigma-Aldrich, A5795), or anti-mouse IgG-HRP (Santa Cruz Biotechnology, sc-2005) secondary antibody. TBST washed membranes were analyzed using the SuperSignal™ West Pico PLUS (Thermo Fisher, 34580) and SuperSignal™ West Femto Maximum Sensitivity Substrate (Thermo Fisher, 34095) using a ChemiDoc™ MP machine (Bio-Rad).

### Yeast Two-Hybrid assay

The lithium acetate method^91^ was used to co-transform respective LexA binding domain (BD) and B42 activation domain (AD) constructs into the yeast strain *EGY4.8 p8op*. Successful transformants were selected by colony growth at 30°C on SD-UHW/Glu (2% (w/v) glucose, 0.139% (w/v) yeast synthetic drop-out medium, pH 6 without uracil, histidine, tryptophan, 0.67% (w/v) BD Difco yeast nitrogen base, 2% (w/v) Bacto agar). Three days post transformation, eight to ten colonies from each sample were combined in 200 µL of water and the concentration of yeast cells was determined, adjusted and diluted to OD_600_ = 1, 0.5, 0.1, 0.01 and 0.001. Ten µL of each dilution was dropped out on SD-UHW/Gal/Raf selection plates to confirm growth (+leucine) and on SD-UHWL/Gal/Raf interaction plates (-leucine). Here, glucose was replaced by 2% galactose and 1% raffinose to enable expression of the LexA BD-fused proteins. Plates were subsequently incubated at 30°C and growth of yeast transformants was documented three days after drop-out unless specified differently.

### Western blot analysis using yeast material

Protein levels of the constructs of interest upon expression in yeast were determined by growing yeast transformants in liquid SD-UHW/Gal/Raf medium at 30°C, 200 rpm for ∼16h. Cells were harvested by centrifugation at 1000 x g for 5 min. For extraction of total protein, yeast cells were collected in 300 µL NH_4_-Acetate buffer (1 M NH_4_-Acetate, 150 mM NaCl, 60 mM Tris-HCl-pH 7.5, 10 mM PMSF, 5 mM EDTA, 2% Protease-Inhibitor-Cocktail (Sigma-Aldrich, P8340) and glass beads (Sigma-Aldrich, G8772) of 300 µL in volume were added. Cells were disrupted using a Retsch Beat Beater (2 min at 30 Hz). After disruption of cells, an additional 500 µL of NH_4_-Acetate buffer was added to each sample and the extract was transferred to a new tube. Samples were incubated on ice for 1.5 h and centrifuged at 16000 x *g* for 10 min. The supernatant was removed and pellets were washed with 1 ml of 1 M NaCl, centrifuged at 16000 x *g* for 10 min and resuspended in 200 µL Urea-SDS buffer (50 mM Tris-HCl pH 6.8, 2% SDS, 8 M urea, 2 mM EDTA, 5% glycerol, 0.004% bromophenole blue, 1% β-mercaptoethanol). Upon an additional centrifugation step at 16000 x *g* for 10 minutes, the proteins from the supernatant were separated by SDS-PAGE (Bio-Rad), and blotted onto polyvinylidene fluoride (PVDF) membranes (Merck). Membranes were blocked in TBST with 3% non-fat milk and probed with either anti-HA (Roche 11867423001 for B42-HA fusion proteins) or anti-LexA (Santa Cruz Biotechnology sc7544 for LexA-fusion proteins) primary antibodies followed by anti-rat (Sigma-Aldrich, A5795) or anti-mouse IgG-HRP (Santa Cruz Biotechnology, sc2005) secondary antibodies as appropriate. TBS-T washed membranes were analyzed using the SuperSignal™ West Pico PLUS (Thermo Fisher, 34580) and SuperSignal™ West Femto Maximum Sensitivity Substrate (Thermo Fisher, 34095) using a ChemiDoc™ MP machine (Bio-Rad).

## Acknowledgements

We thank Nicolaus von Wirén and Ricardo Giehl for useful discussion on the role of plant iron homeostasis during pathogen attack; Petra Bauer for suggestions on iron supplementation experiments; Volker Lipka for information on the preparation of chitin for ROS measurements; the UoC Institute for Plant Sciences’ greenhouse team for their expertise in maintenance and high-yield propagation of barley; Dennis Schapelhouman and Saskia Bodendorf for maintaining *Bh* isolate K1; Paul Schulze-Lefert for making the MPIPZ facilities available for the generation of transgenic *AVR_a13_* lines; Anne Harzen for MS sample preparation; Gabriel Hacker, Sabine Haigis and Emma Crean for technical assistance in propagating barley plants and *in vitro* growth of barley under differential iron conditions, generating *AVR_a_*-overexpressing lines, and the selection transgenic barley lines, respectively. This work was funded by the Deutsche Forschungsgemeinschaft (DFG, German Research Foundation) Emmy Noether Programme (SA 4093/1-1 to IMLS), the DFG’s Collaborative Research Centre Grant (SFB-1403—414786233 to IMLS, GD), the Germany’s Excellence Strategy CEPLAS (EXC-2048/1, project 390686111 IMLS, GD, GH) and the Max-Planck-Society (HN).

## Data availability

Source Data are provided with this paper. The mass spectrometry proteomics data have been deposited to the ProteomeXchange Consortium via the PRIDE partner repository with the dataset identifier PXD071751.

## Author contributions

Experiments: WS (proximity labelling, Y2H, CoIP, Split-LUC, confocal microscopy, pathogen infection, selection of *srf3* barley), MB-S (Y2H, cell death assays, confocal microscopy, pathogen infection, ROS), SCStolze (LC-MS/MS), MPP (root growth inhibition), ISD and SCSent (protoplast assay), PC and GH (barley *srf3* line design, generation and recovery), MH and IMLS (generation, selection, assessment and propagation of transgenic plants, generation of T2 and T3 transgenic lines). Data analysis: WS, MB-S, SCStolze, IMLS. Investigation: WS, MB-S, GD, HN, IMLS. Supervision: GH, HN, IMLS. Funding acquisition: IMLS, GD. Project administration, conceptualization and study design: IMLS. Writing: WS, MBS, IMLS with input from all authors. Editing: All authors.

