## Supplementary Information for "Targeting of the barley cell-surface receptor SRF3 by the *Blumeria hordei* effector AVRA13 overlaps with AVRA13 recognition by MLA and the induction of NLR-mediated cell death"

**Supplementary Data 1:** Liquid Chromatography-Mass Spectrometry analysis of proteins identified by proximity-dependent protein labelling in wild-type and transgenic barley lines expressing *BirA* and *$AVR_a$-BirA* constructs. Standard analysis refers to LFQ data-based enrichment and volcano plot analysis refers to iBAQ data-based enrichment. Proteins enriched in respective samples are highlighted by orange color. Bait proteins are highlighted by green color.

**Supplementary Fig. 1: Enrichment of $AVR_A$-interacting proteins by BioID-based biotin labelling. a** Schematic representation of the BioID-based proximity labelling strategy used to isolate proteins in proximity to $AVR_A$ effectors in barley. **b** Increase of biotin treatment beyond 250 µM does not lead to increased biotinylation of barley proteins by $AVR_{A1}$-BirA-4myc. Leaf tissue from wild type Golden Promise (GP)Fast and stable transgenic barley expressing *$AVR_{a1}$-BirA-4myc* was treated with 0, 100, 250 and 600 µM biotin for 36 hours. Total protein was extracted and biotinylation was determined using streptavidin-HRP (Strep-HRP) western blot. **c, e, g** Comparison of $AVR_{A1}$-BirA-4myc and $AVR_{A13}$-BirA-4myc (c), $AVR_{A13}$-BirA-4myc and $AVR_{A10}$-BirA-4myc (e) and $AVR_{A7}$-2-BirA-4myc and $AVR_{A9}$-BirA-4myc (g) protein levels in respective stable transgenic barley across three replicates (R1, R2, R3) as determined by anti-myc western blot; wild type GP Fast and BirA-4myc lines were included as controls throughout. **d, f, h** Strep-HRP western blot of biotinylated proteins upon streptavidin-based enrichment in wild type GP Fast, BirA-4myc and respective $AVR_A$-BirA-4myc samples upon treatment of leaf tissue with 250 µM biotin. Samples represent one of the biological replicates subjected to LC-MS/MS analysis for protein identification.

**Supplementary Fig. 2: Analysis of $AVR_{A}$ interactors. a** ShinyGO-based gene ontology (GO) of proteins identified specifically as interactors of **$AVR_{A1}$**, **$AVR_{A7}$**, **$AVR_{A9}$**, **$AVR_{A10}$** and **$AVR_{A13}$** by proximity-dependent protein labelling (Fig. 1a). Analysis was conducted using ShinyGO v0.77 (https://bioinformatics.sdstate.edu/go77/) with a false discovery rate (FDR) cut-off of < 0.05, revealing the potential biological roles or molecular activities of the host proteins enriched upon proximity-dependent protein labelling involving $AVR_{A}$-BirA-4myc lines.

**Supplementary Fig. 3: a** Phylogenetic trees (Maximum Likelihood method) and domain structure of *Arabidopsis thaliana* and barley Strubbelig Receptor Family (SRF) members. **b** Phylogenetic tree (Neighbor-Joining method) of SRFs.

**Supplementary Fig. 4: a** Sequence alignment of barley HvSRF3-like proteins. (HvSRF3-L1, HvSRF3-L2, HvSRF3-L3, and HvSRF3-L4). Domains of HvSRF3 are highlighted with colored boxes. SUB: Strubbelig domain, LRR: Leucine-rich repeat domain, JM: Juxtamembrane domain. Conserved residues are shaded in dark black, and similar residues are shaded in light black. The alignment highlights the conserved regions and sequence diversity among the different isoforms. **b**. AlphaFold3-based structural prediction of the cytoplasmic domain from the HvSRF3-like proteins. Blue: kinase domains (pTMs >80), orange: juxtamembrane domains (pTMs <70).

**Supplementary Fig. 5: a** Protein levels of assays displayed in Fig. 2c. Leaf material was harvested three days post *Agrobacterium*-mediated transformation of leaves from four-week-old *N. benthamiana* plants. Total protein was extracted and detected using anti-HA and anti-myc western blot (WB) analyses. CBB: Coomassie brilliant blue staining of respective membranes post WB detection. **b and c** *Nicotiana benthamiana* leaves were transiently transformed with the constructs indicated. At three days post transformation, protein-protein interaction was determined by the complementation of the luciferase (LUC) reporter by nLUC and cLUC-fused protein upon addition of substrate (b) and GFP-TRAP-based immunoprecipitation of GFP or mYFP tagged proteins followed by anti-GFP and anti-HA western blot analysis (c). Figures shown are representatives of at least two independent experiments. **d** Protein levels of assays displayed in Fig. 2e. Yeast transformants were grown in selective medium containing leucine to $OD_{600}$=1. Cells were harvested for total protein extraction. Proteins were separated by gel electrophoresis and detected using anti-LexA-based (LexA-BD, receptor domains) or anti-HA-based (HA-B42-AD, GFP or effectors) western blot analyses as indicated. CBB: Coomassie brilliant blue staining of respective membranes post WB detection.

**Supplementary Fig. 6: a, b, d, e** Protein levels of assays displayed in Fig. 3. Yeast transformants were grown in selective medium containing leucine to $OD_{600}$=1. Cells were harvested for total protein extraction. Proteins were separated by gel electrophoresis and detected using anti-LexA-based (LexA-BD, receptor domains) or anti-HA-based (HA-B42-AD, GFP or effectors) western blot analyses as indicated. CBB: Coomassie brilliant blue staining of respective membranes post WB detection. **c** Yeast growth upon $AVR_{A13}$ interaction assays with HvSRF3-L1, HvSRF3-L2, HvSRF3-L3 and HvSRF3-L4. Direct comparison of interaction intensities of figures displayed in Fig. 2e and Fig. 3b. **f** AlphaFold models of HvSRF3-$L3^{KD}$ (pTM =0.9) and HvSRF3-$L4^{KD}$ (pTM =0.88). The N-lobe, C-lobe, P-loop, DFG motif, activation loop, catalytic loop of respective residues in HvSRF3-$L3^{KD}$ and $L4^{KD}$ are coloured cyan, pale green, orange, blue, red and yellow, respectively.

**Supplementary Fig. 7: a** Protein levels corresponding assays shown in Fig. 4a. Yeast transformants were grown on selective medium containing leucine to $OD_{600}$=1. Cells were harvested for total protein extraction. Proteins were separated by gel electrophoresis and detected with anti-LexA (LexA-BD) or anti-HA antibodies (B42-AD) by western blot analyses as indicated. **b, d and f** Protein levels corresponding assays shown in Fig. 4d and 4e (b), Fig. 4f and 4g (d) and 4h-4j (f). Leaf material was harvested three days post *Agrobacterium*-mediated transformation of leaves from four-week old *Nicotiana benthamiana* plants. Total protein was extracted and detected using anti-myc-based, anti-LUC-based, anti-GFP-based or anti-mCherry-based western blot (WB) analyses as indicated. CBB: Coomassie brilliant blue staining of respective membranes upon WB detection. **c** Fluorescent signal of mCherry or $AVR_{A13}$-mCherry corresponding to split-YFP assay shown in Fig. 4f. **e** Representative close up of split-YFP assays involving HvSRF3 and HvBAK1. *N. benthamiana* leaves were transiently transformed with constructs encoding mCherry or $AVR_{A13}$-mCherry together with C-terminally $YFP^{N}$-fused HvSRF3 and C-terminally $YFP^{C}$-fused HvBAK1 constructs as indicated. **g and h** *N. benthamiana* leaves were transiently transformed with constructs as indicated. At two days post transformation, mCherry or YFP fluorescence signal was detected using a Leica TCS SP8 confocal laser scanning microscope (g) or proteins were extracted and separated by gel electrophoresis and detected with anti-HA or anti-mCherry antibodies by western blot analyses as indicated (h). CBB: Coomassie brilliant blue staining of respective membranes post WB detection. Pictures shown are representative results from at least two independent biological replicates. Scale bars: 25 µm. Signal peptides were depleted from all effector-encoding constructs throughout.

**Supplementary Fig. 8:** PAMP-mediated production of reactive oxygen species (ROS) in leaves of barley transgenic lines expressing *BirA* or *AVR$_{a13}$-BirA*. Chitin- and flg22-mediated production of reactive oxygen species (ROS) in leaves of barley transgenic lines expressing *BirA* or *AVR$_{a13}$-BirA*. ROS production was elicited by the addition of **a and b** 10 µg/ml chitin or **c and d** 100 nM flg22 as indicated and luminescence was determined upon addition of peroxidase and luminol-based substrate in a Tecan plate reader. Results are depicted as relative luminescence unit (RLU) per minute for 30 min of measurement. Values were obtained from at least two independent experiments. Differences between treatments were assessed by ANOVA ($p = 0.0778$ for b, $p = 0.07642$ for d). n.s. = not significant.

**Supplementary Fig. 9: a** Protein levels correspond to yeast two-hybrid assays displayed in Fig. 5. Yeast transformants were grown on selective medium containing leucine to $OD_{600}$=1. Cells were harvested for total protein extraction. Total proteins were separated by gel electrophoresis and detected with anti-LexA (LexA-BD) or anti-HA antibodies (B42-AD) by western blot analyses as indicated. **b** Protein levels correspond to split-luciferase assays displayed in Fig. 5. Leaf material was harvested three days post *Agrobacterium*-mediated transformation of leaves from four-week old *Nicotiana benthamiana* plants. Total protein was extracted and detected using anti-myc-based or anti-HA-based western blot analyses as indicated. **c** AVR$_{A13}$ protein detection in *Arabidopsis thaliana* wild type Col-0 and FLAG-*AVR$_{a13}$* lines as determined by anti-FLAG western blot analysis. CBB: Coomassie brilliant blue.

**Supplementary Fig. 10: a** *Blumeria hordei* (*Bh*) infection rate on barley plants grown *in vitro* under varying iron levels (10µM, 100 µM and 250 µM FeNaEDTA) as determined by *Bh* biomass quantification. **b and c** *Bh* disease symptoms on leaves of independent transgenic barley lines expressing *BirA* or *AVR$_{a9}$-BirA* at five days post infection upon treatment with tween (mock) or 0.5 g/L FeEDDHA. Figures shown are representatives of at least three independent experiments (b). Quantification of *Bh* colonies (c) was performed from pictures taken of the adaxial and abaxial site of the infected leaves across all replicates of experiments described in b. Differences between treatments were assessed by Kruskal–Wallis test ($p<2.2e-16$) and subsequent Dunn's test. Samples marked by identical letters in the plots do not differ significantly ($p>0.05$) in the Dunn's test. **d** Schematic overview of CRISPR-Cas9-generated mutant alleles of barley HvSRF3 variants indicating position of guide RNA (gRNA) target sites. **e** *srf3* mutant lines under standard growth conditions. **f to i** PAMP-mediated production of reactive oxygen species (ROS) in leaves of barley wild-type and *srf3* mutant lines. ROS production was elicited by the addition of 10 µg/ml chitin (f and g) or 100 nM flg22 (h and i) as indicated and luminescence was determined upon addition of peroxidase and luminol-based substrate in a Tecan plate reader. Results are depicted as relative luminescence unit (RLU) per minute for 30 min of measurement. Values were obtained from at least three independent experiments. Differences between treatments were assessed by ANOVA ($p = 0.61$ for g, $p = 0.32$ for i). n.s. = not significant.

**Supplementary Fig. 11: Protein levels corresponding to yeast two-hybrid and cell death assays displayed in Fig. 7. a, b and d** Yeast transformants were grown in selective medium containing leucine to $OD_{600}$=1. Cells were harvested for total protein extraction. Total proteins were separated by gel electrophoresis and detected with anti-LexA (LexA-BD) or anti-HA antibodies (B42-AD) by western blot analyses as indicated. **c** Leaf material was harvested three days post *Agrobacterium*-mediated transformation of leaves from four-week-old *Nicotiana benthamiana* plants. Total protein was extracted and detected using anti-GFP-based western blot (WB) analyses. CBB: Coomassie brilliant blue staining of respective membranes post WB detection.

**Supplementary Fig. 12: a** *Nicotiana benthamiana* leaves were transiently transformed with constructs encoding HvSRF3-mCherry together with constructs encoding GFP, $\mathrm{AVR}_{\mathrm{A13}}$ wild type or its mutants and mCherry fluorescence signals were determined using a Leica TCS SP8 confocal laser scanning microscope two days post transformation. Pictures shown are representative results from at least three independent biological replicates. Scale bars: 25 µm. **b** Quantification of mCherry signals across all replicates of experiments described in a. **c** Quantification of GFP/YFP signals across all replicates of experiments described in a. Differences between samples were assessed by Kruskal–Wallis (p= 2.2e-16 for I and p= 0.003206 for c) and subsequent Dunn's test. Samples marked by identical letters in the plots do not differ significantly (p>0.001) in the Dunn's test. **d** Mesophyll protoplasts of barley cultivar Golden Promise wild-type and *srf3* lines were co-transfected with a *pZmUBQ:luciferase* reporter plasmid, *Mla13,* and constructs carrying either *GFP,* the $AVR_{a1}$ control or $AVR_{a13}$ all in *pIPKb002*. Luciferase activity was determined at 16 h post transfection and normalized to the respective *GFP* sample. Values were obtained from at least four independent transfections performed on at least three independent days. Differences between samples were assessed by Kruskal–Wallis (p= 8.06E-08) and subsequent Dunn's test. Samples marked by identical letters in the plots do not differ significantly (p>0.05) in the Dunn's test. Signal peptides were depleted from all effector-encoding constructs throughout.

### Supplementary Fig. 1

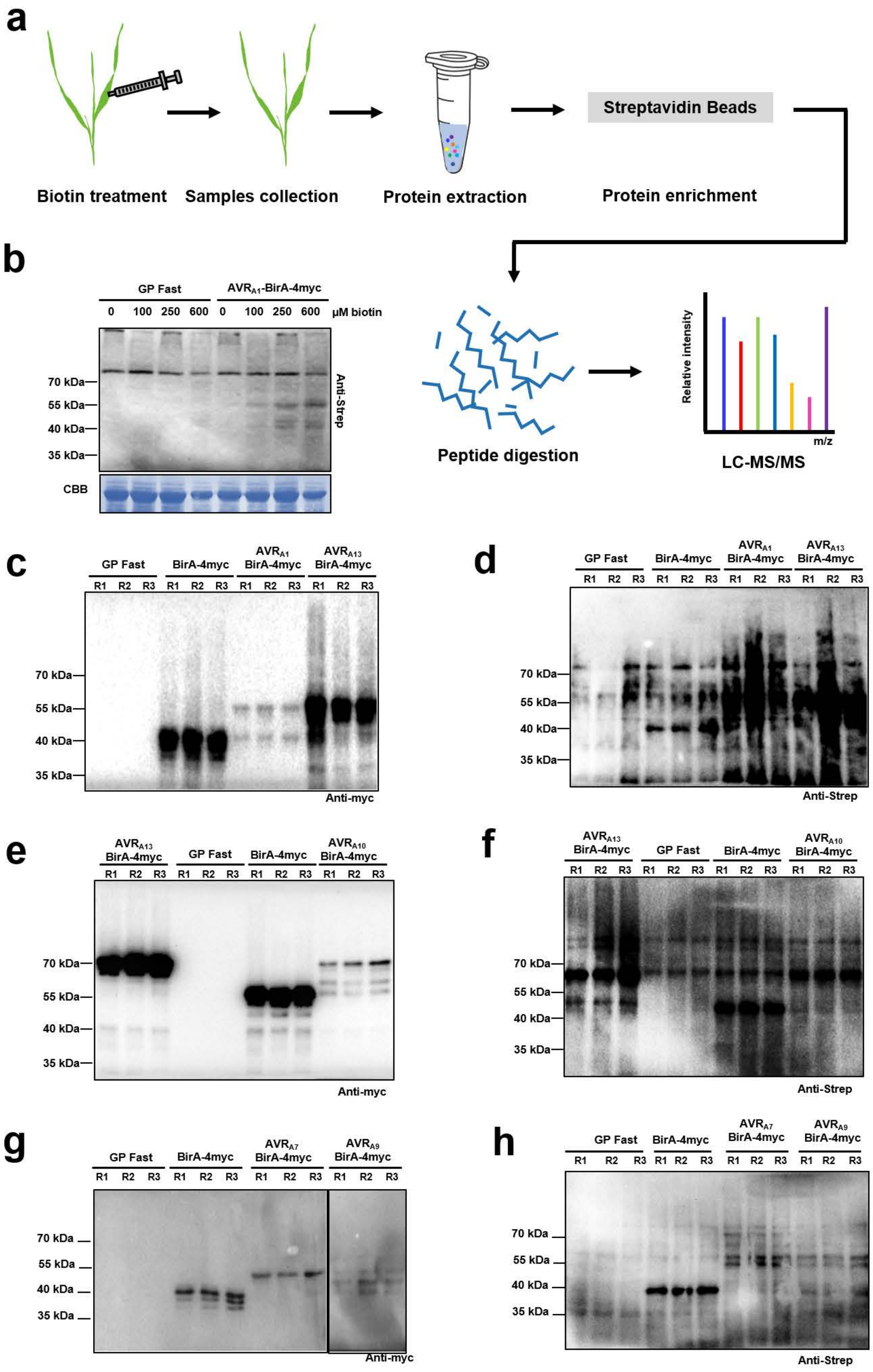

### Supplementary Fig. 2

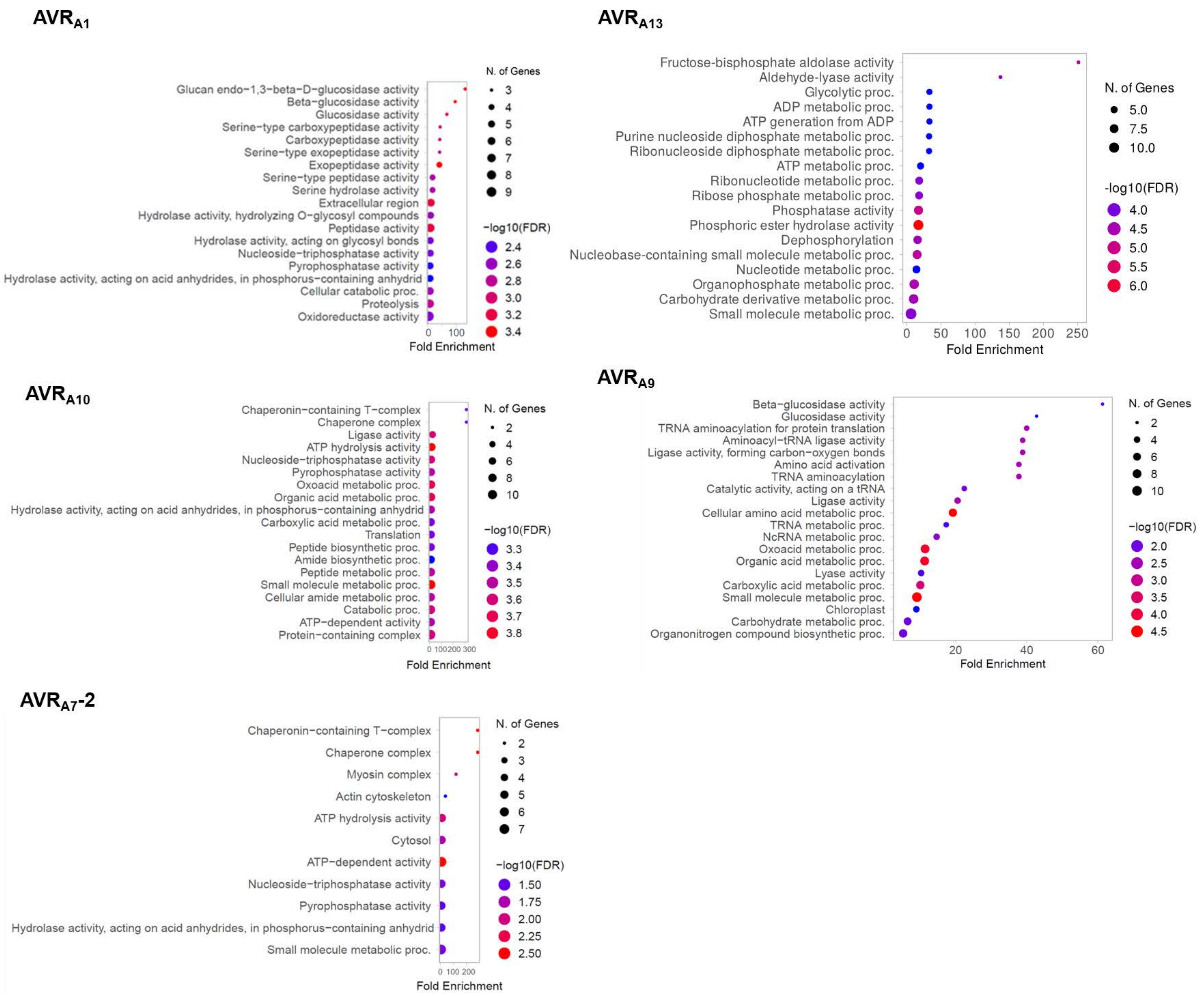

### Supplementary Fig. 3

**a**

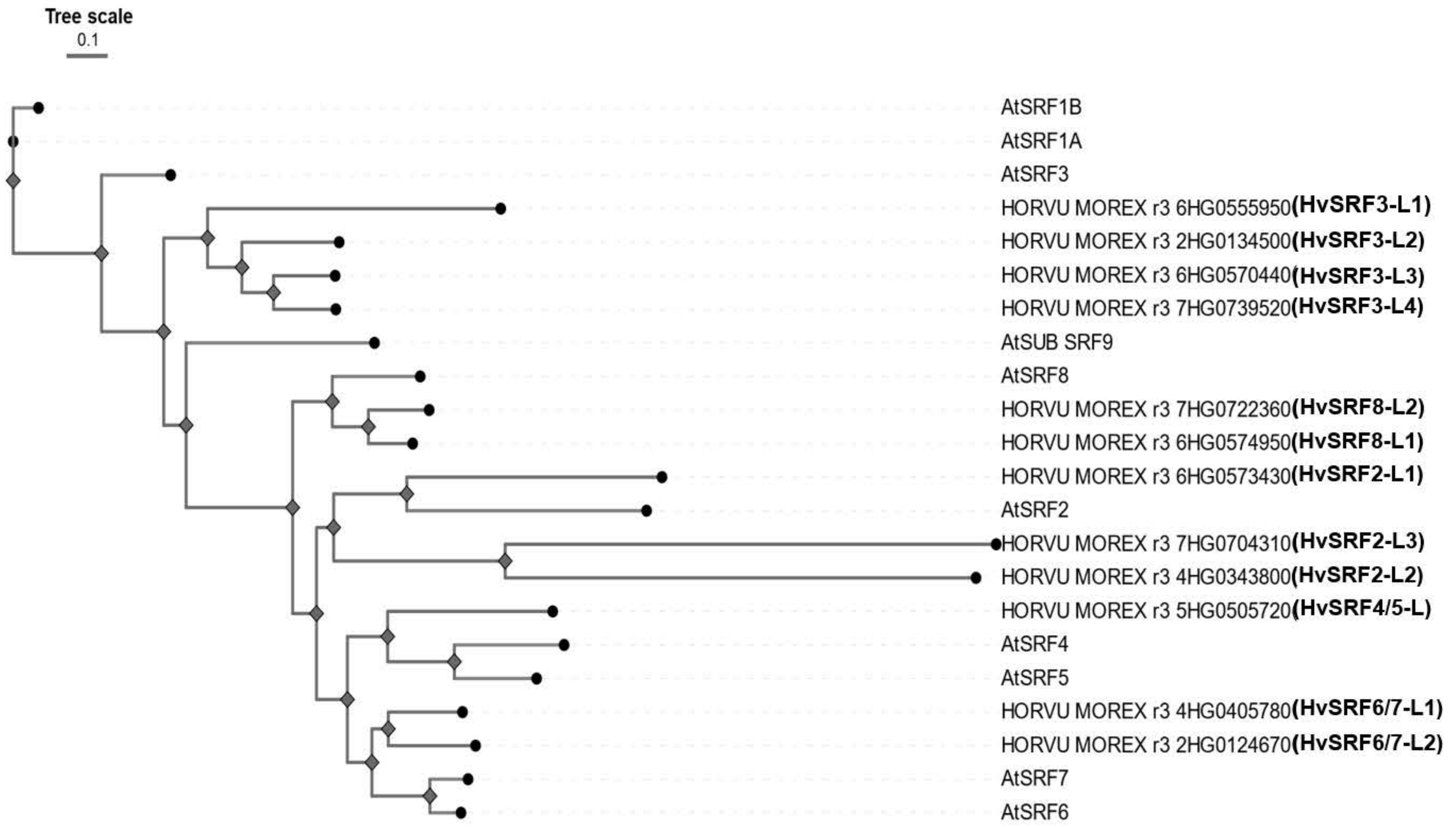


**b**

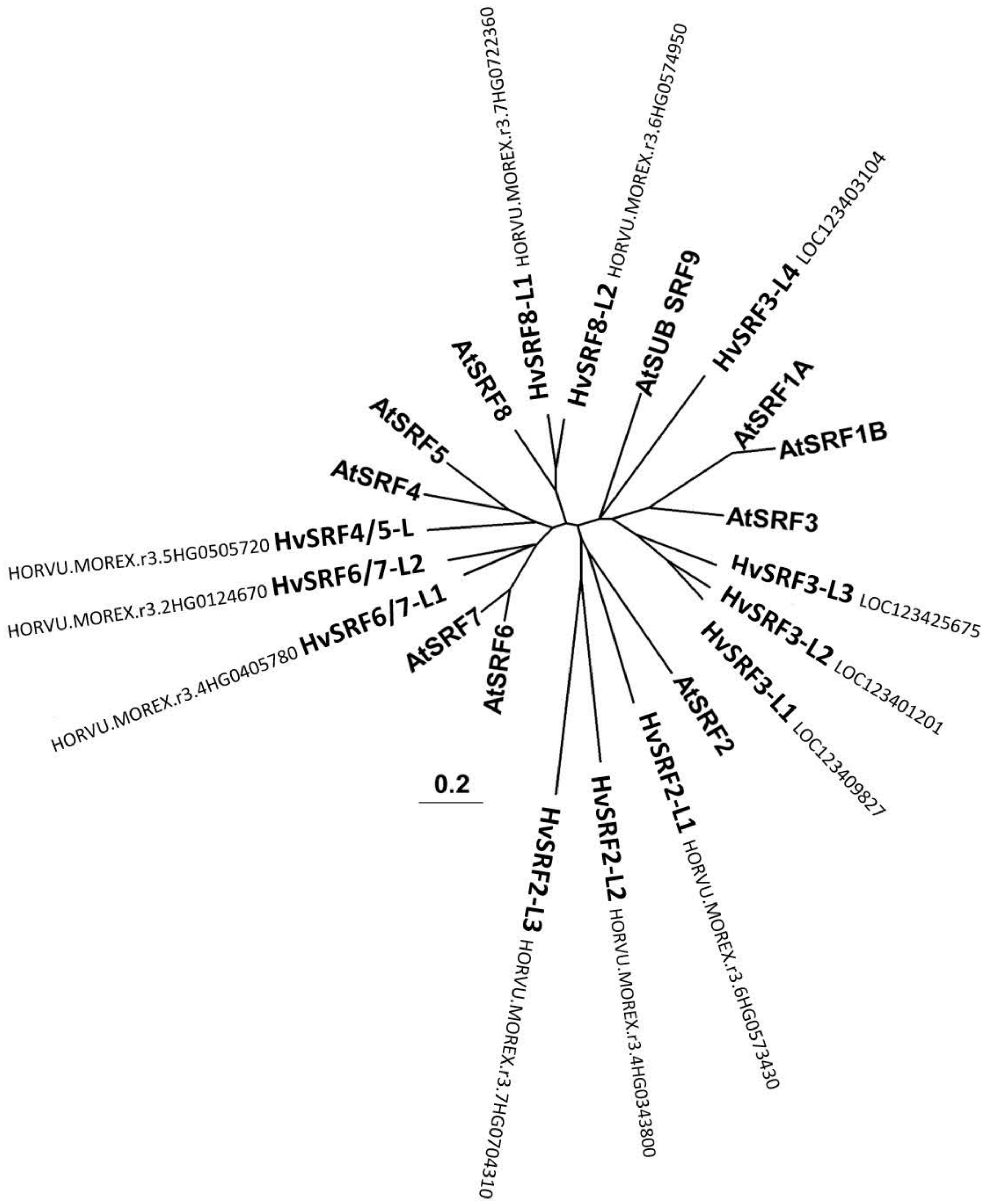

#### Supplementary Fig. 4

**a**

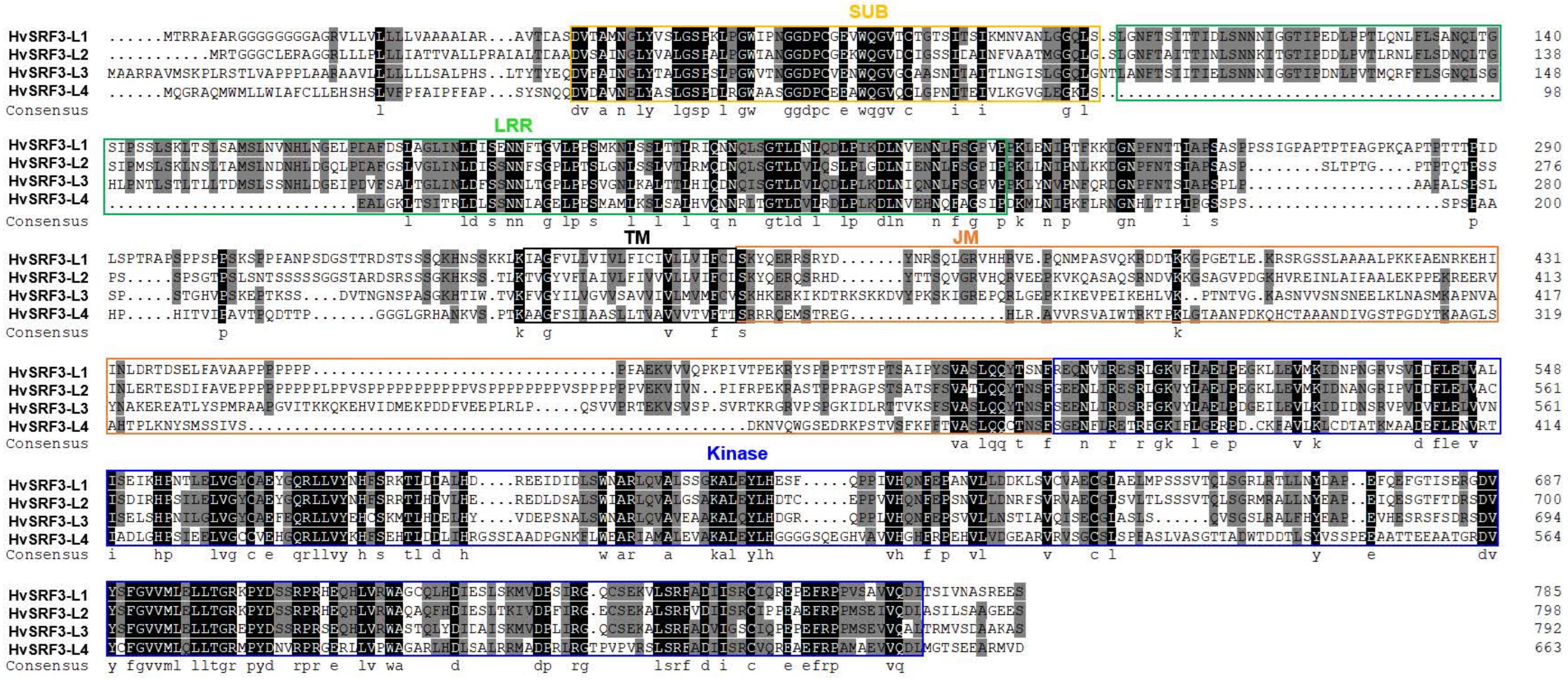


**b**

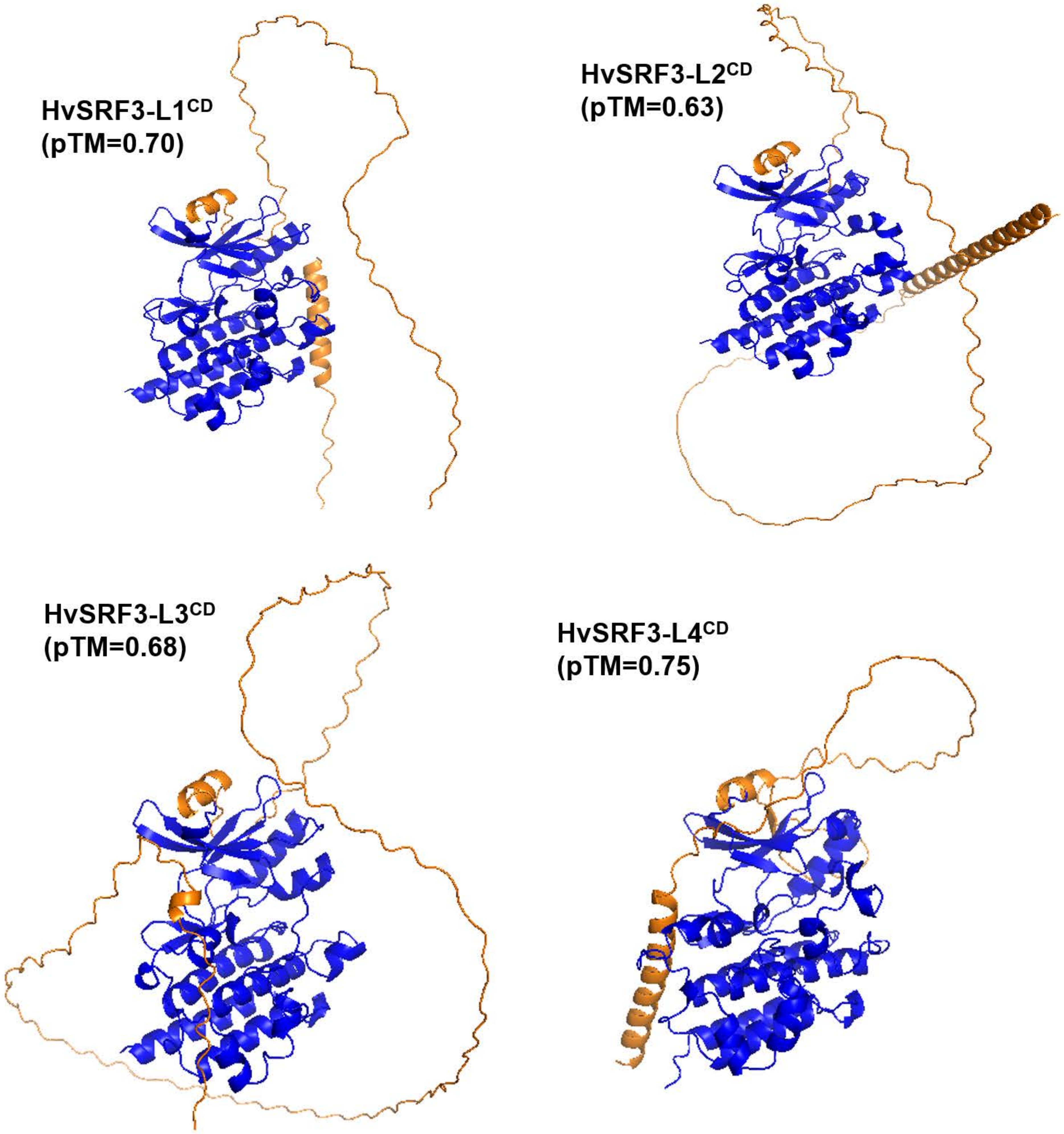

### Supplementary Fig. 5

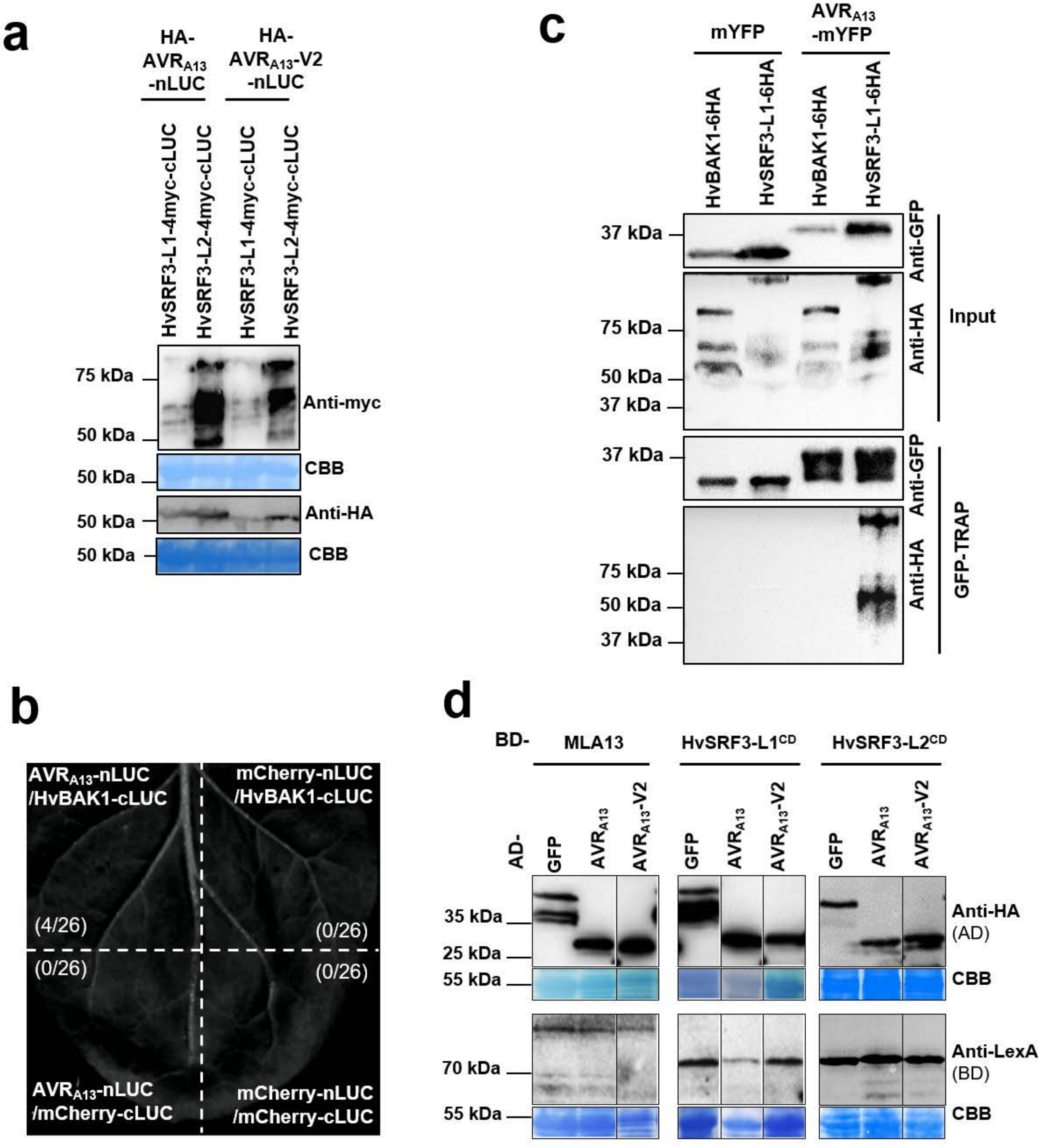

#### Supplementary Fig. 6

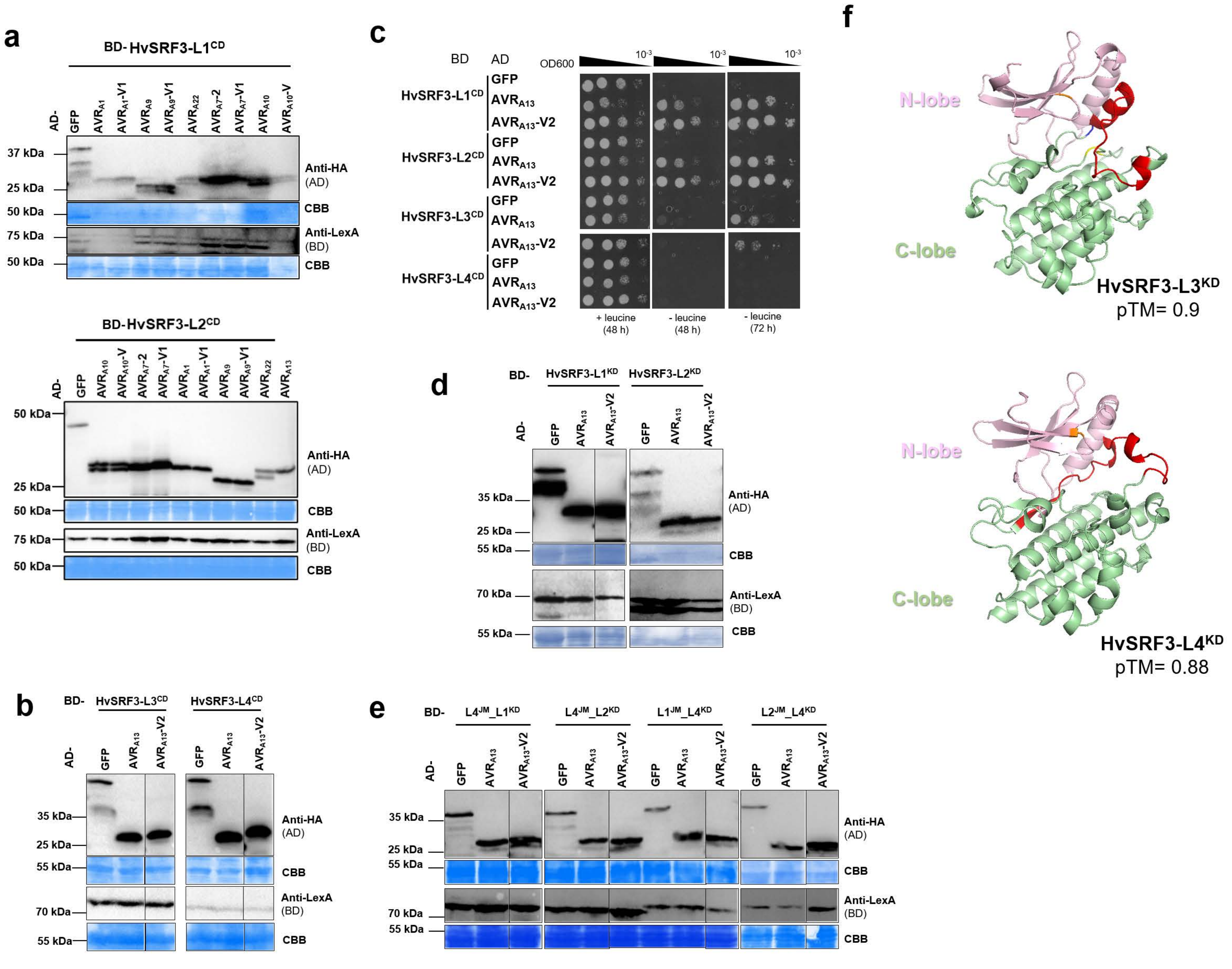

### Supplementary Fig. 7

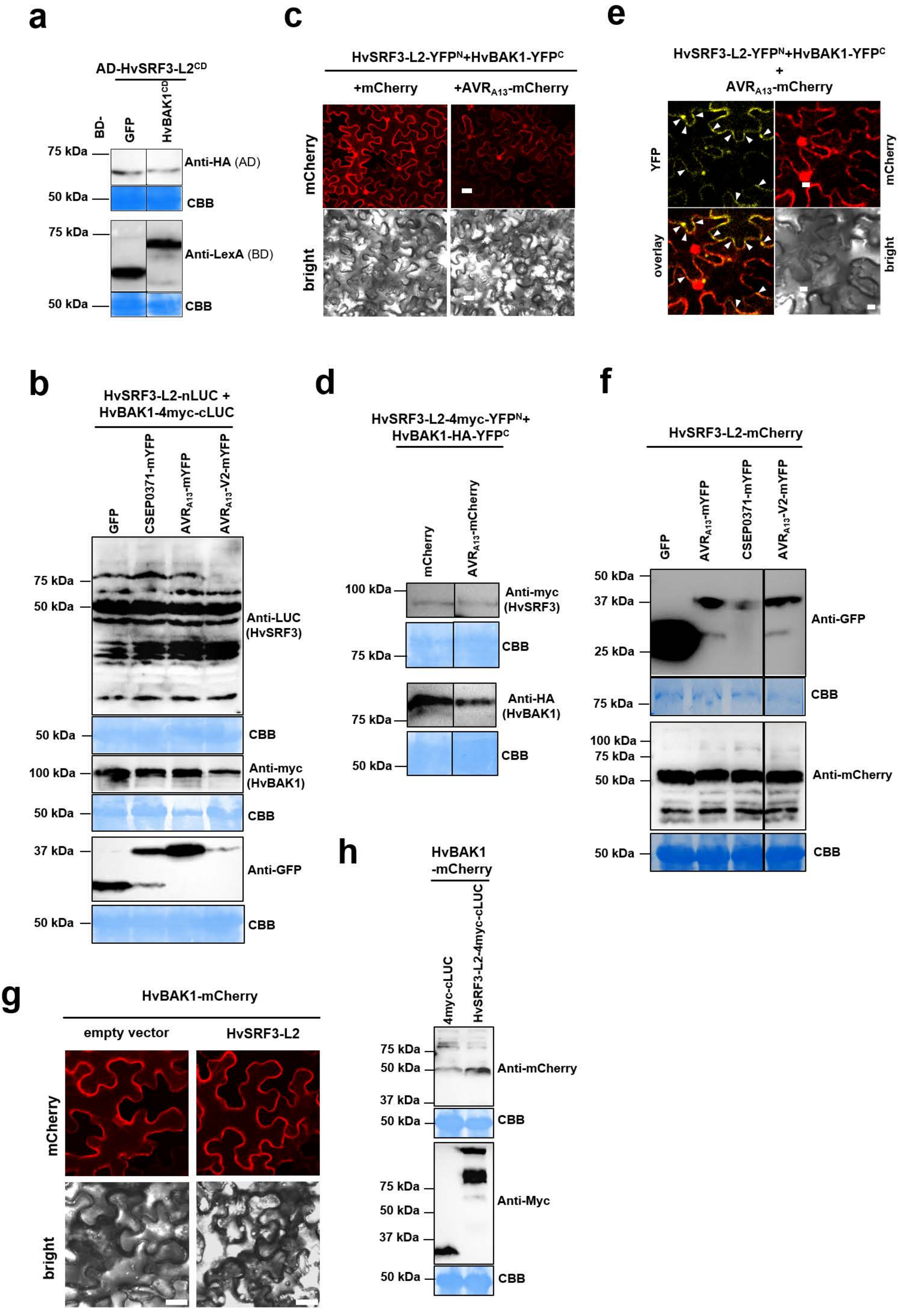

### Supplementary Fig. 8

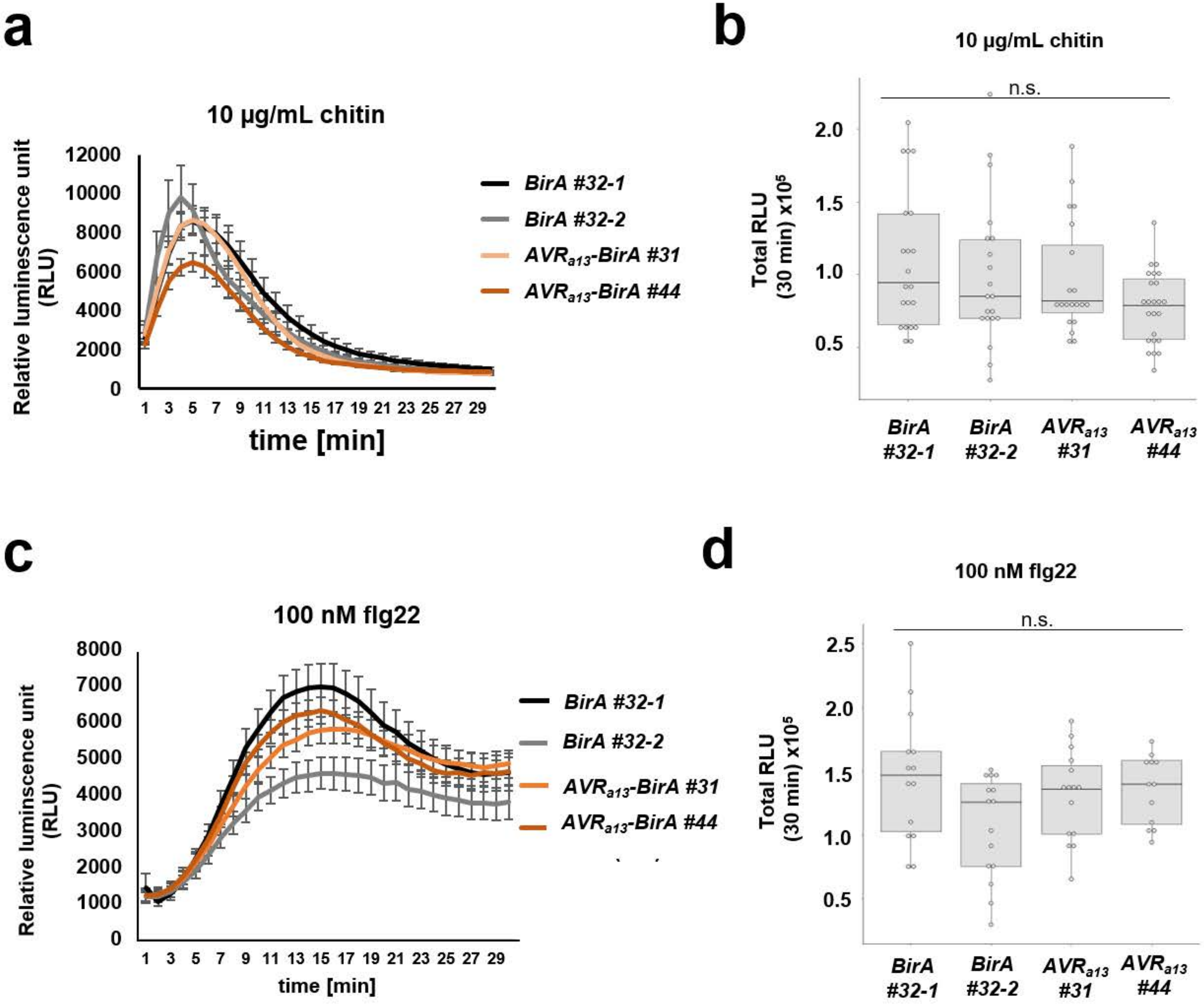

### Supplementary Fig. 9

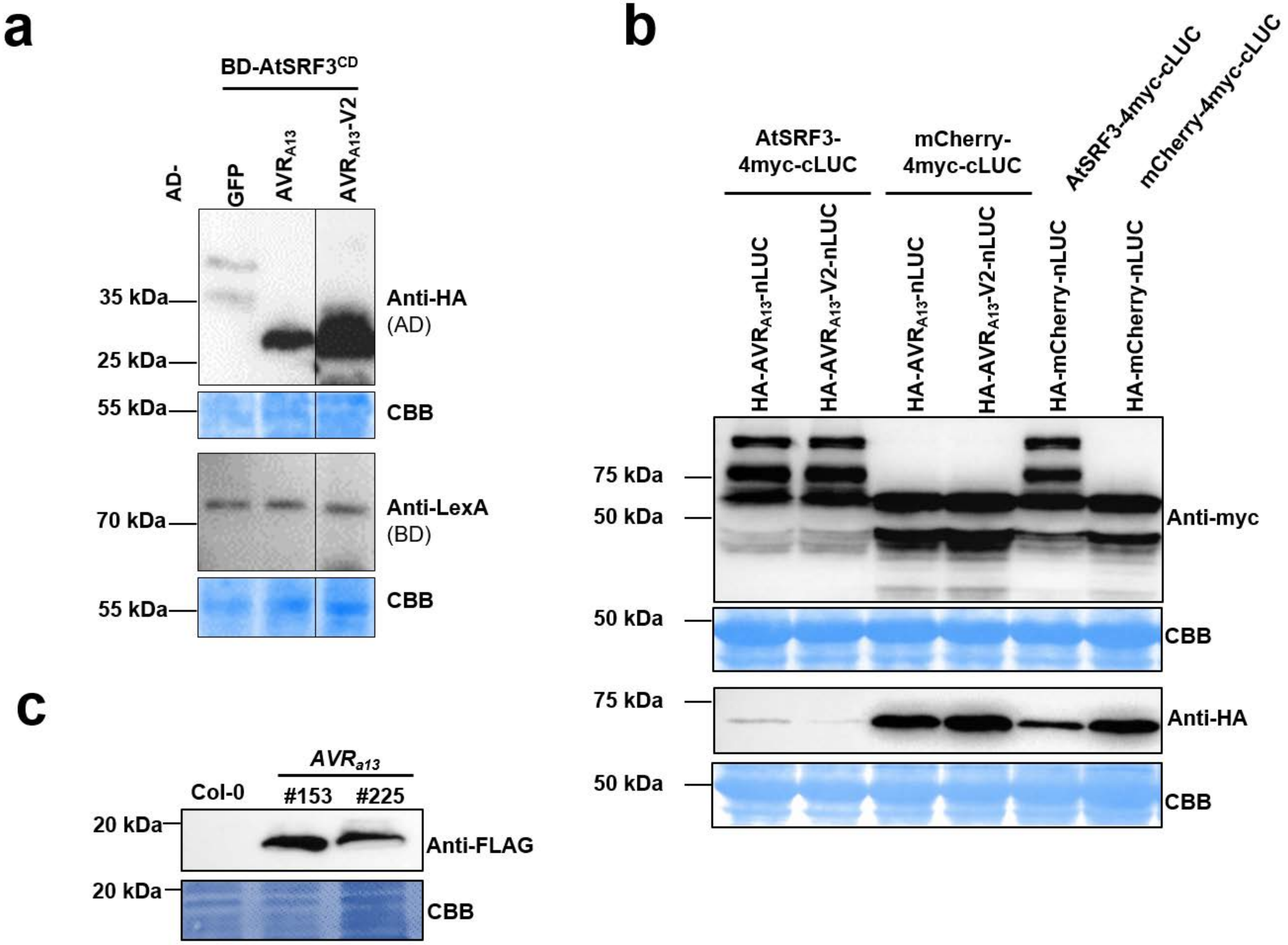

### Supplementary Fig.10

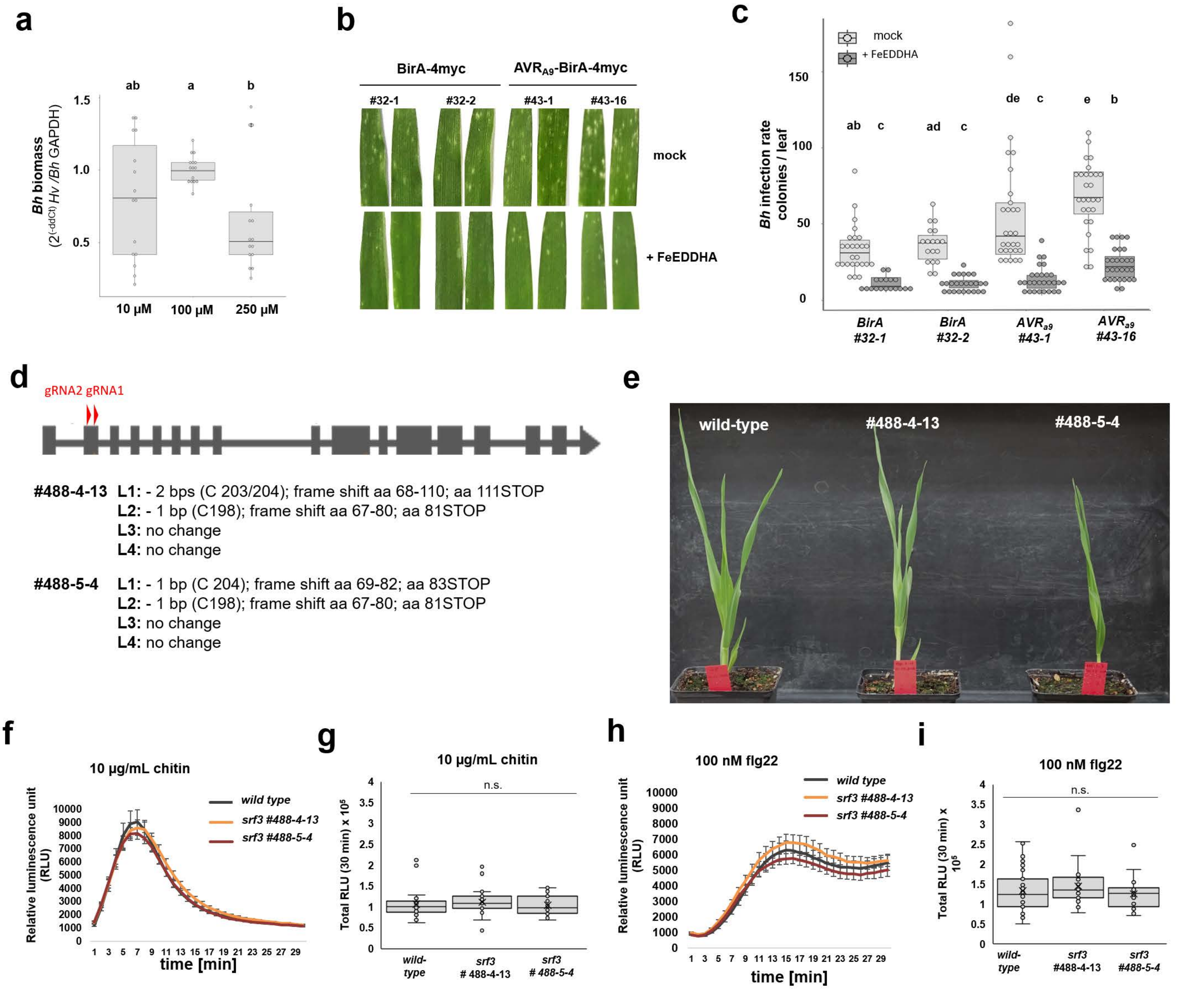

### Supplementary Fig. 11

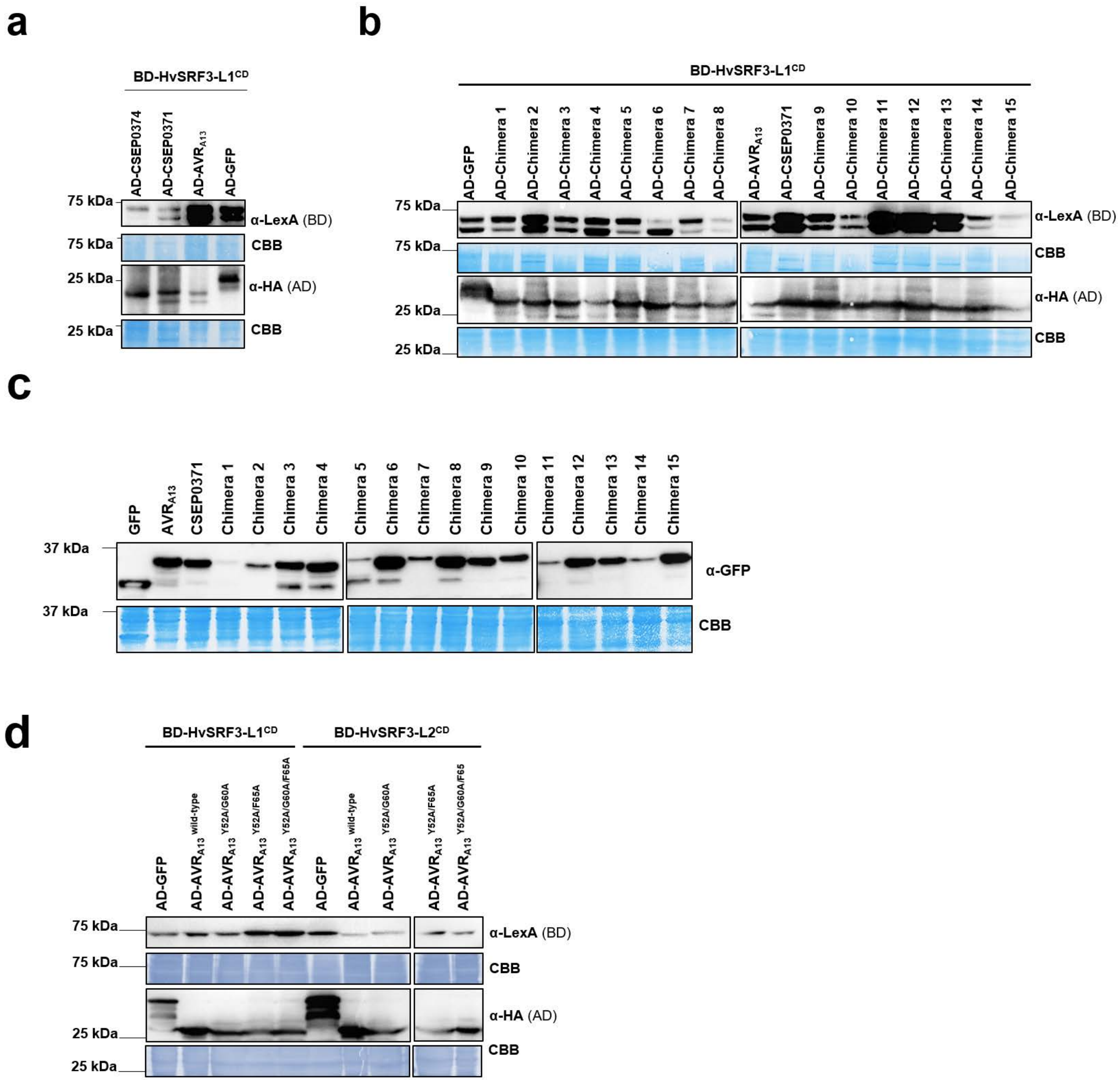

### Supplementary Fig. 12

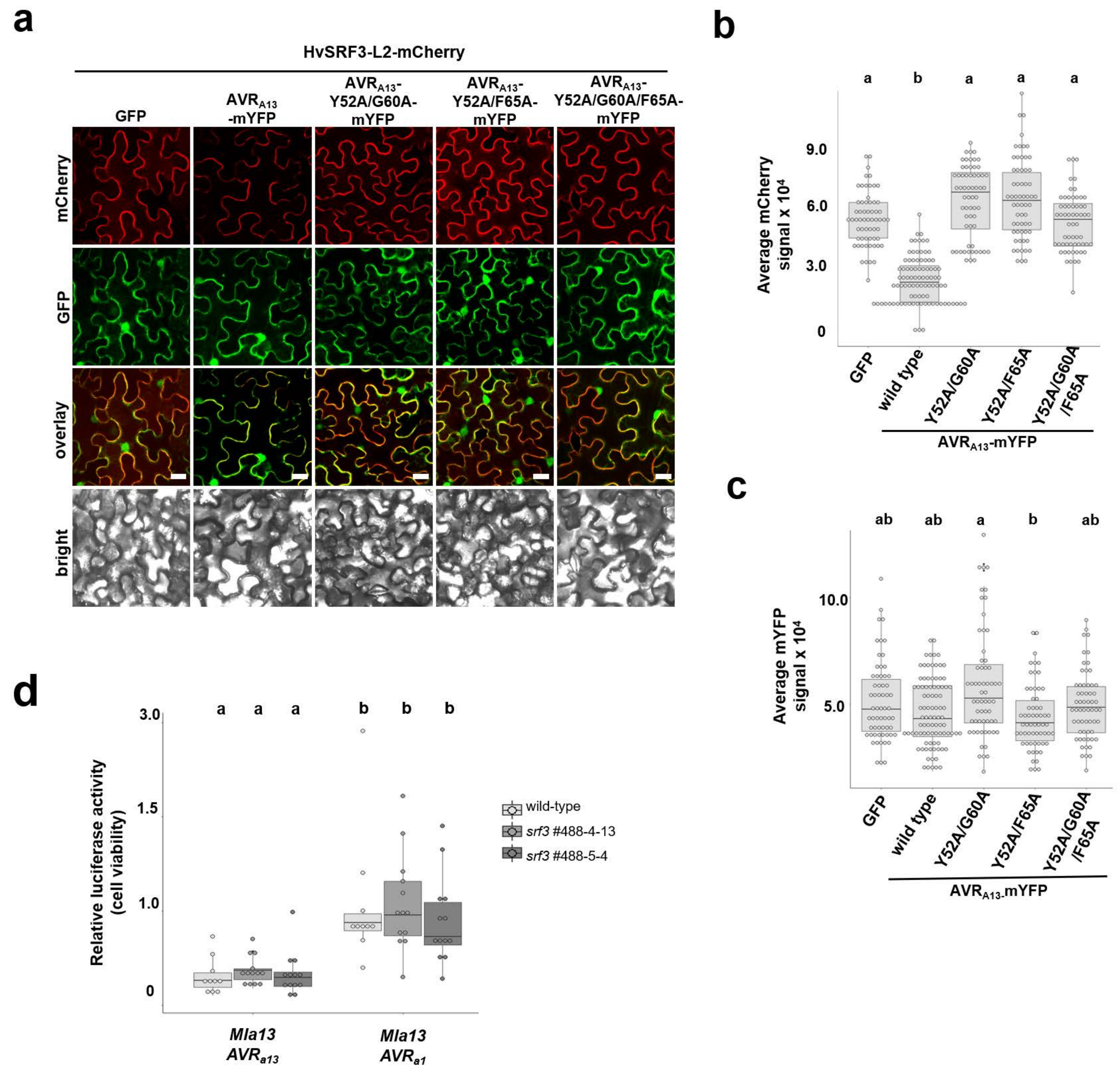
